# Rising, falling, stalling: Complex biodiversity trajectories in UK rivers

**DOI:** 10.1101/2025.11.21.689659

**Authors:** Mansi Mungee, Martin Wilkes, Vicky Bell, William E. Kunin, Christopher Hassall, Lee E. Brown

## Abstract

Freshwater ecosystems face intense anthropogenic pressures, with global indicators showing steep biodiversity declines in many regions. Yet in parts of Europe, including the UK, several recent analyses of long-term monitoring datasets have documented widespread improvements in macroinvertebrate communities following major water-quality investments. These gains are often interpreted as being nationally uniform, based on spatially aggregated data or simplified effect sizes, despite substantial site-level heterogeneity in temporal dynamics. Consequently, the prevalence, distribution, and causes of local biodiversity dynamics remain poorly resolved. We address this gap using macroinvertebrate time-series data from 803 river sites in England’s Environment Agency monitoring network. We applied an unsupervised, shape-explicit time-series clustering framework to classify sites into distinct shape-based clusters for three complementary metrics: pollution sensitivity of communities (Whalley, Hawkes, Paisley & Trigg (WHPT) Index), total abundance, and Simpson’s diversity index. We then characterized taxonomic and environmental differences among clusters using Linear Discriminant Analysis and non-metric multidimensional scaling. Finally, we quantified how temporal trajectories are structured across increasingly aggregated spatial scales (site, water body, catchment, typology, region) using a Bayesian variance-partitioning model, providing the first multi-metric, multi-scale assessment of macroinvertebrate trends in UK rivers. Overall, our analyses reveal substantial heterogeneity in ecological trajectories: although many sites showed improving trends, more than half (∼53%) were in decline or lacked clear directional change. Sites with consistent increases or declines were strongly differentiated by longitudinal position, channel morphology, and water chemistry, indicating that network placement and local environmental context mediate river biodiversity changes. Variance partitioning showed that most explainable variation lies at the site and typology levels, demonstrating that broader regional or national summaries can obscure a mosaic of non-linear, site-level trajectories and persistent local declines. Collectively, these findings affirm that freshwater biodiversity change in UK rivers cannot be summarized by a national or even regional narratives of recovery, but represents a mosaic of asynchronous, site-specific trajectories. These results illustrate the need for a renewed investment in dense, long-term river monitoring networks capable of resolving biodiversity change across hierarchical spatial scales.

## Introduction

Freshwaters are among the most threatened ecosystems on the planet, and headline figures of 85% declines in indicator species abundance (WWF, 2024) have led to numerous calls to ‘bend the curve’ by changing the way rivers are managed and restored (e.g. Tickner et al., 2020). Aquatic organisms are particularly sensitive to environmental change due to their diverse biological and ecological traits (Wilkes et al., 2020), the spatial isolation of rivers in landscapes which limit efficient dispersal (Carrara et al., 2012; Heino et al., 2015; Tonkin et al., 2018; Carraro et al., 2023), and because hydrological pathways act as corridors or sinks for upstream stressor inputs (Vannote et al., 1980; Pringle, 2003; Freeman et al., 2007). Together, these factors limit the occurrence of refugia or buffer zones for organisms against environmental stressors (Revenga et al., 2005; Dudgeon et al., 2006; Reid et al., 2019). Yet, against this global backdrop of concern, studies from several European countries, including the UK, have reported widespread gains in freshwater macroinvertebrate richness, and, in some cases, abundance (Vaughan and Ormerod, 2014; Van Klink et al., 2020; Vaughan and Gotelli, 2019; Pharaoh et al., 2023).

Regional improvements in water quality across Europe have followed investments in sewage treatment (e.g. the Urban Waste Water Treatment Directive 91/271/EEC, EU Water Framework Directive 2000/60/EC; European Commission, 1991; European Commission, 2000; Whelan et al., 2022; Johnson et al., 2019), better treatment of pollution from industries such as mining (e.g. ∼232 billion litres treated annually by 82 schemes in the UK in 2024–25; Mining Remediation Authority, 2025), and/or recovery from acidification (Murphy et al., 2014; Medupin, 2020). However, some pollutants, including road runoff and new tyre chemicals, may be getting worse, and nutrient and pesticide runoff appear to be increasing over time as agricultural intensification continues (Wagner et al., 2018; Halsband and Sørensen, 2020; Schürings et al., 2024). At the same time, legacy pressures have not disappeared and long-standing nutrient issues persist in many systems (e.g. increase in chlorophyll-a despite declines in phosphorus; Worrall et al., 2026). Against this backdrop, the extent and generality of “good news” stories remain contested because biodiversity estimates are highly sensitive to data processing choices (harmonisation of taxonomy, choice of diversity metrics, and handling of abundance classes), trend-model specifications (sign-only vs. non-linear shape), and the spatial scale at which data are aggregated, all of which vary across studies (Demars, 2024; Wilkes et al., 2025a; Wilkes et al., 2025b). Understanding which river systems are either improving or declining, and why this is the case, is vital to developing robust recovery plans for global freshwater biodiversity.

In the UK specifically, the magnitude and even the direction of change differ markedly across studies based upon the same national dataset (Vaughan and Ormerod, 2012, 2014; Murphy et al., 2014; Outhwaite et al., 2020; Qu et al., 2023; Powell et al., 2023). Very few works have compared temporal variation at individual sites (e.g. Vaughan and Ormerod, 2012, 2014; Qu et al., 2023), but the end results are typically summarised as directional classes (either as single values of slopes, or effect sizes for numbers of increase/decrease/no change) rather than explicit trajectory shapes. Yet, management-relevant features of both recovery and decline are often complex and non-linear, with features such as point step-changes after major pollutant incidents or restoration interventions (Johnson et al., 2019), episodic responses to extreme events (Fornaroli et al., 2020), temporal lags in biodiversity response across taxonomic and functional richness (Pharaoh et al., 2023), and potential tipping points, regime shifts or alternative states (Carrier-Belleau et al., 2023; Hernández Martínez de la Riva et al., 2023). Methods that reduce trajectories to a single slope or even a sign miss these dynamics, overlooking complex, intervention-sensitive behaviour across individual sites.

National or regional scale trends provide scientists and environmental managers with important ‘big picture’ estimates of change (e.g. Outhwaite et al., 2020; Powell et al., 2023; Pharaoh et al., 2023; Johnson et al., 2025). However, mechanistically, the presence of site-level variation in biodiversity responses is a consequence of interactions among multiple physical, chemical and biological processes which vary at local scales (Murphy et al., 2014; Civan et al., 2018; Verberk et al., 2016). For instance, recent studies show that the ecological imprint of water chemistry is inherently local and can propagate non-uniformly downstream, often outperforming models built on land use or temperature alone (Fornaroli et al., 2020; Nguyen et al., 2024; Johnson et al., 2025). Impacts of large-scale drivers such as climate warming can also vary across sites depending on local pollution legacies or flow regime (Durance and Ormerod, 2007; Vaughan and Gotelli, 2019). Biological drivers include dispersal events, density-dependent processes, and ecological drift (Thompson et al., 2020). Site-level responses can thus depart strongly from catchment or regional means, and national trends can flatten local recovery or decline. Yet, systematic attention to site-level variation in river biological trend studies remains limited.

Cross-scale analyses present an opportunity to uncover the mechanisms linking local stressors to large-scale biodiversity patterns (Pharaoh et al., 2023; Powell et al., 2023). However, no previous study has assessed the variation in freshwater biodiversity responses across the full hierarchy from from individual sites, through water bodies and catchments, to broader typology classes and regions, nor have they compared complex, non-linear, site-level time series across multiple metrics in a single, hierarchically nested and spatially explicit framework (but see Vercelloni et al., 2014 for a marine ecosystem application). In this study, we address these two important gaps: (1) we identify and delineate clusters of sites that exhibit distinct non-linear temporal trajectories with respect to pollution sensitivity (with the widely used Whalley, Hawkes, Paisley and Trigg (WHPT Total) index; Paisley et al., 2014), total macroinvertebrate abundance, and taxonomic diversity, and (2) use variance partitioning across the nested hierarchy to evaluate site-level patterns in the context of broader catchment and regional characteristics.

We tested the following hypotheses: (H1) macroinvertebrate communities will not exhibit simple monotonic increases or decreases; instead, we expected clusters of sites with varied non-linear, temporal shapes that reflect asynchrony in multiple environmental drivers; (H2) environmental drivers will strongly differentiate cluster membership at the individual-site level, consistent with site-specific chemistry and/or hydromorphology; (H3) taxonomic composition would differ between clusters with increasing, decreasing, or stable trajectories, with declines in WHPT associated with the loss of high-scoring families that are sensitive to stressors, and increases linked to their persistence or recovery; and (H4) the largest share of explainable variation across clusters will sit at the site level, supporting the importance of local drivers and cautioning against over-interpretation of regional or national averages. We harmonised taxonomy (family level), derived explicit shape-based time-series clusters for each metric, and then analysed the environmental and compositional correlates across the clusters. Finally, we used a hierarchical Bayesian variance decomposition analysis to quantify how cluster assignment varies across the (nested) spatial scales. Our study provides the first multi-metric, cross-hierarchy analysis of river ecosystem biodiversity trends and addresses the ecological granularity at which stressors act.

## Materials & Methods

### Macroinvertebrate Data

Macroinvertebrate data were sourced from the Environment Agency’s (EA) long-term monitoring surveys (https://environment.data.gov.uk/ecology/explorer/). Synonymous taxa were updated to accepted names, abundances aggregated to familial level, and we retained samples from 2002–2023, when the EA adopted more accurate abundance estimates, improved quality control, and standardised taxa lists (Wilkes et al., 2025a). Data were filtered to: (i) 3-minute kick-sampling protocol (Murray-Bligh, 1999); (ii) abundances on a linear scale or converted from log scale; (iii) sites sampled in spring for ≥10 of 20 years, with one spring (March - May) sample randomly selected if multiple were collected per season; (iv) sites with data in at least one of 2002–2004 and one of 2021–2023; and (v) sites without missing samples for ≥3 consecutive years. To ensure comparability across biodiversity metrics, we retained only WHPT scoring taxa for all analyses.

### Statistical Analysis

We generated site-specific time series for WHPT Total (abundance-based adaptation of the globally utilised Biological Monitoring Working Party score system; Chesters, 1980; Hawkes, 1998), total abundance, and Simpson diversity index, representing pollution sensitivity, productivity, and diversity, respectively. We opted for WHPT Total rather than WHPT-ASPT (Average Score Per Taxon); while ASPT attempts to correct for sampling effort and seasonal effects, in our dataset, these effects were standardised by restricting analyses to a single sample during spring. We recognise, however, that residual variation (e.g. collector differences, subtle changes in sampling conditions or identification) may still affect trends (Jones et al., 2023). To minimise these potential biases, we restricted our analyses to samples collected after 2002, when the Environment Agency implemented stricter quality control procedures, including standardised taxonomic lists and strengthened QA/QC through systematic analyst competency testing, ring tests, reverse ring tests, and random re-checking of samples by a second analyst (Murray-Bligh, 1999; Environment Agency, 2014, 2017). Metrics were range-normalised (between 0 and 1) and missing values interpolated using the function *na_interpolation* from the package imputeTS v3.3 (Moritz et al., 2017) in the R programming environment (R Core Team, 2024).

### Time-Series Clustering

Time-series clustering partitions ‘*n’* time series into ‘*k’* clusters (*k << n*) such that statistically similar series are grouped together based on some similarity/distance measure (Aghabozorgi et al., 2015). Several methods are available for unsupervised clustering of time series. Here, we used nine commonly used approaches, combining three clustering methods (hierarchical, partitioning k-means, and fuzzy c-means) with three distance measures: Euclidean, Dynamic Time Warping (DTW), and shape-based distance (SBD) (Pavoine, 2009; Bloomfield et al., 2018; Wei et al., 2020; Hegg and Kennedy, 2021; see **Supporting Information S1** for details on each of these). Since the challenges of clustering complex time-series data are not unique to ecology, analytical approaches from other disciplines may be applicable, including measures to evaluate cluster validity and robustness (Ranacher and Tzavella, 2014). Ecological data, however, are typically more heterogeneous, often contain gaps, and rarely span the long temporal scales needed to assess clustering efficiency. In this study, datasets covered ∼20 years, compared to climatic or financial series that often extend up to 100 years (Sathiaraj et al., 2019; Thomas et al., 2021). To address these limitations, we simulated additional data with statistical properties similar to the empirical macroinvertebrate series. This enabled us to first optimise models and evaluate clustering efficiency on simulated datasets, and then use the ‘best-fitting’ model for the observed, empirical data. We simulated a large time-series dataset with statistical properties similar to the empirical macroinvertebrate time series using autoregressive integrated moving average (ARIMA) model parameters (**Supporting Information S1**). This dataset was used to compare the different methods, evaluate clustering efficiency, and determine the optimal number of clusters (k) (Kaufman and Rousseeuw, 1990; Arbelaitz et al., 2013; Kim and Ramakrishna, 2005) using a suite of internal Cluster Validity Indices (CVIs; Aghabozorgi et al., 2015; **see Supporting Information S2 for more information on CVIs**). Prior to clustering, time series were smoothed with Nadaraya–Watson kernel regression using Gaussian GCV-optimised bandwidths (λ) to reduce inter-annual noise (Nadaraya, 1964; Watson, 1964). Clustering was implemented with the function *tsclust* from the package dtwclust v5.5.12 (Sarda-Espinosa, 2017; Sarda-Espinosa et al., 2018) for all three metrics of WHPT, abundance, and diversity.

### Linear Discriminant Analysis (LDA)

We used Linear Discriminant Analysis (LDA) to assess the contribution of different environmental factors in distinguishing the time-series clusters. We assembled a suite of river morphological and water chemistry variables (**Table 1**; also see **Supporting Information S3**). All predictor variables were mean centred and standardised to unit variance before analysis, and Variance Inflation Factor (VIF) analysis was performed to subset to the least correlated variables with independent axes of information. We used 10-fold cross-validation, where in each iteration 90% of the data were used for training and 10% for testing. LDA was performed using the *lda* function from the package MASS v7.3-60.0.1 (Ripley et al., 2013). Model evaluation and goodness of fit were evaluated using a confusion matrix (caret v7.0-1; Kuhn, 2008; Kuhn et al., 2020) and Cohen’s Kappa values (K). Performance was assessed using three standard measures: sensitivity, specificity, and accuracy, using both overall (across all clusters) and class-specific (per cluster) statistics. ELSA (entropy-based local indicator of spatial association) statistics were calculated to evaluate neighbourhood spatial autocorrelation in the time-series clusters (package elsa v1.1-28; Naimi et al., 2019) and tested for any global spatial structure using enterograms (Naimi et al., 2019).

**Table 1.**
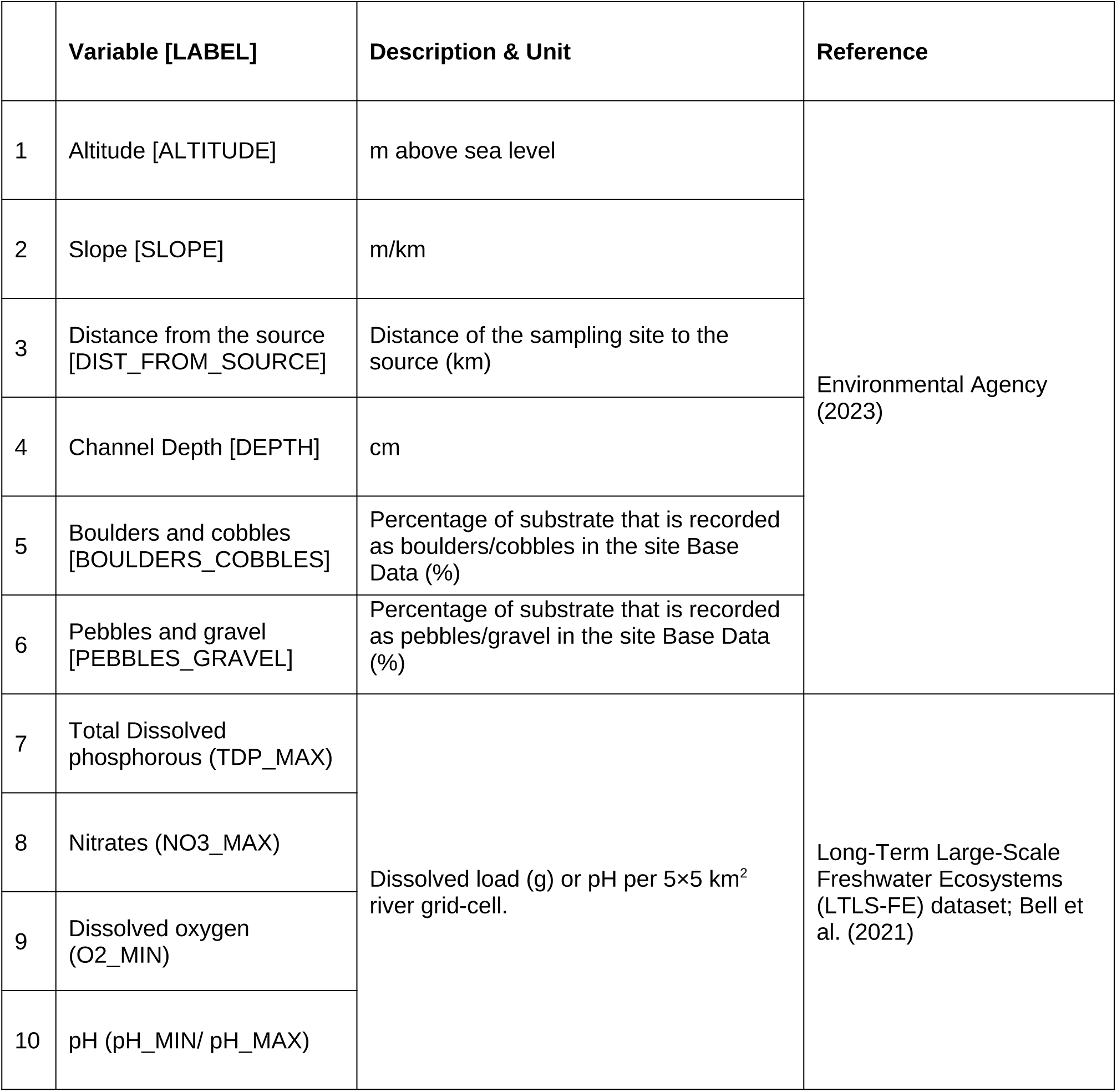
Environmental variables used in Linear Discriminant Analysis (LDA) of time-series cluster membership. Variables represent a combination of geomorphological attributes (altitude, slope, distance from source, channel depth, and substrate composition) derived from Environment Agency datasets, and water chemistry indicators (total dissolved phosphorus, nitrates, dissolved oxygen, and pH) obtained from the Long-Term Large-Scale Integrated Model (LTLS-IM; Bell et al., 2021). All variables were standardized (mean-centred and scaled to unit variance) prior to analysis, and highly collinear predictors were excluded following Variance Inflation Factor (VIF) analysis to ensure independent axes of information. Only the final set of independent variables used in LDA are shown(For the full (initial) list see Supporting Information S3).

|  | Variable [LABEL] | Description & Unit | Reference |
| --- | --- | --- | --- |
| 1 | Altitude [ALTITUDE] | m above sea level | Environmental Agency (2023) |
| 2 | Slope [SLOPE] | m/km |  |
| 3 | Distance from the source [DIST_FROM_SOURCE] | Distance of the sampling site to the source (km) |  |
| 4 | Channel Depth [DEPTH] | cm |  |
| 5 | Boulders and cobbles [BOULDERS_COBBLES] | Percentage of substrate that is recorded as boulders/cobbles in the site Base Data (%) |  |
| 6 | Pebbles and gravel [PEBBLES_GRAVEL] | Percentage of substrate that is recorded as pebbles/gravel in the site Base Data (%) |  |
| 7 | Total Dissolved phosphorous (TDP_MAX) | Dissolved load (g) or pH per 5×5 km<sup>2</sup> river grid-cell. | Long-Term Large-Scale Freshwater Ecosystems (LTLS-FE) dataset; Bell et al. (2021) |
| 8 | Nitrates (NO3_MAX) |  |  |
| 9 | Dissolved oxygen (O2_MIN) |  |  |
| 10 | pH (pH_MIN/ pH_MAX) |  |  |

### Non-metric multidimensional scaling (NMDS)

To explore the taxonomic composition of clusters, we used non-metric multidimensional scaling with 100 iterations (NMDS; function *metaMDS* from vegan v2.6-8; Oksanen et al., 2007). We calculated distance matrices using Bray–Curtis distance on a transformed (square-root) community data frame (Cluster ID × Taxa). Distance-based permutational multivariate analysis of variance (PERMANOVA; Anderson, 2014), which compares the variation between groups to the variation within groups, was performed to test if familial compositions differ significantly across the different time-series clusters for each metric (WHPT, abundance, and diversity). PERMANOVA was implemented with the function *adonis2 (*vegan v2.6-8; Oksanen et al., 2007).

### Hierarchical variance decomposition

We fit an intercept-only variance-components model using a Bayesian mixed-effects (hierarchical) approach in the brms v2.22.0 package (Bürkner, 2017, 2021). Our aim was to estimate how much variability in freshwater macroinvertebrate responses is attributable to each level of the monitoring hierarchy:

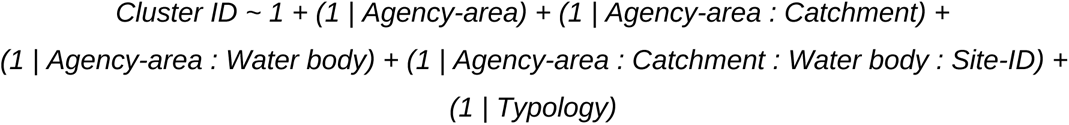

where the agency area represents the geographical administrative divisions used by the Environment Agency in England. Thus, catchments and water bodies are treated as parallel units within each agency area, and sites are nested within the intersection of agency, catchment, and water body. In addition, we included Typology (System A typology; Directive, 2000; **Table 2**; also see **Supporting Information S4**) as a cross-classified grouping factor, recognising that river types defined by geology, altitude, and catchment size cut across administrative and hydrological boundaries. The categorical response variable, Cluster_ID, represents groupings of sites with similar temporal trajectories in WHPT, abundance, or diversity. For each metric, the baseline category was set to the cluster that did not show either a strong increasing or decreasing trend. A multinomial distribution with a logit link was chosen to model these categorical outcomes. We deliberately used an intercept-only model because our focus was not on estimating the effects of predictors (e.g. time or covariates) on cluster membership, but rather on partitioning the variance in site clustering across multiple hierarchical and cross-linked levels. By excluding fixed predictors, the model attributes variability entirely to the random effects, yielding a direct estimate of the relative contribution of each spatial scale (agency region, typology, catchment, water body, site) to the observed clustering patterns (for a semi-parametric application with fixed effects, see Vercelloni et al., 2014). We used four MCMC chains (parallelised across four cores) with 4,000 iterations each (2,000 warm-up), setting adapt_delta = 0.95 and max_treedepth = 12 to improve sampling efficiency and ensure convergence. Convergence was assessed through graphical posterior predictive checks (function *pp_check*), the potential scale reduction factor (R̂), and effective sample sizes (Bulk-ESS and Tail-ESS).

**Table 2.** River typology classification used in this study (UK Technical Advisory Group; UKTAG Typology for Rivers 2003). System A Typology (UKTAG 2003), which was used in this study, classifies rivers according to fixed physical and chemical descriptors (altitude band, catchment size, and dominant geology) yielding 18 discrete “GB Type” categories. System B simplifies this framework by grouping sites into six broader classes defined by geology (Calcareous, Siliceous, Organic) and altitude (Low < 200 m, High ≥ 200 m). The table lists each System A type, its descriptive category, the corresponding System B class, and the number of monitoring sites from this study that fall within each type.

| Altitude band (m) | Catchment size band (km<sup>2</sup>) | Dominant Geology | System A Typology<sup>†</sup> | Description | System B Typology | Number of sites in this study |
| --- | --- | --- | --- | --- | --- | --- |
| <200 | 10–100 | Siliceous | I | Lowland, small, siliceous streams | VI (Siliceous Low) | 84 |
| <200 | 10–100 | Calcareous | II | Lowland, small, calcareous streams | II (Calcareous Low) | 373 |
| <200 | 10–100 | Organic | III | Lowland, small, organic streams | IV (Organic Low) | 4 |
| <200 | 100–1000 | Siliceous | IV | Lowland, medium, siliceous rivers | VI (Siliceous Low) | 43 |
| <200 | 100–1000 | Calcareous | V | Lowland, medium, calcareous rivers | II (Calcareous Low) | 228 |
| <200 | 100–1000 | Organic | VI | Lowland, medium, organic rivers | IV (Organic Low) | 2 |
| <200 | >1000 | Siliceous | VII<sup>‡</sup> | Lowland, large, siliceous rivers<sup>‡</sup> | VI (Siliceous Low) | 3 |
| <200 | >1000 | Calcareous | VIII | Lowland, large, calcareous rivers | II (Calcareous Low) | 32 |
| <200 | >1000 | Organic | IX | Lowland, large, organic rivers | IV (Organic Low) | 0 |
| 200–800 | 10–100 | Siliceous | X | Mid-altitude, small, siliceous streams | V (Siliceous High) | 22 |
| 200–800 | 10–100 | Calcareous | XI | Mid-altitude, small, calcareous streams | I (Calcareous High) | 4 |
| 200–800 | 10–100 | Organic | XII | Mid-altitude, small, organic streams | III (Organic High) | 4 |
| 200–800 | 100–1000 | Siliceous | XIII | Mid-altitude, medium, siliceous rivers | V (Siliceous High) | 1 |
| 200–800 | 100–1000 | Calcareous | XIV | Mid-altitude, medium, calcareous rivers | I (Calcareous High) | 2 |
| 200–800 | 100–1000 | Organic | XV | Mid-altitude, medium, organic rivers | III (Organic High) | 0 |
| 200–800 | >1000 | Siliceous | XVI | Mid-altitude, large, siliceous rivers | V (Siliceous High) | 0 |
| 200–800 | >1000 | Calcareous | XVII | Mid-altitude, large, calcareous rivers | I (Calcareous High) | 0 |
| >800 | 10–100 | Siliceous | XVIII | High-altitude, small, siliceous streams | V (Siliceous High) | 0 |

<sup>†</sup>Great Britain (GB) Type codes follow UKTAG 2003 Work Program – Task 2a. Typology for Rivers; TAG 2003 WP 2a (03) Rivers typology (Pv3-4-02-04) <sup>‡</sup> Type 7 was noted by UKTAG as rare and requiring validation.

All computations were performed using the R statistical software (version 4.4.2; R Core Team, 2024), on an Ubuntu platform (24.04) with x86_64-pc-linux-gnu.

## Results

The Environment Agency dataset comprised 107,937 macroinvertebrate samples from 15,585 sites collected between 2002 and 2023. After applying the filtering criteria, 27,667 samples from 803 sites were retained for analysis. These sites were distributed across 19 agency areas, 13 WFD typologies, 240 catchments, and 479 water bodies, and represented a wide variation across all environmental and morphological attributes (**Fig. S1 & S2**).

In the simulated time-series dataset, fuzzy Cluster Validity Indices (CVIs) showed inconsistent patterns across the different clustering approaches: 2 out of 5 indices were highest for the SBD distance measure, whereas the other 3 were higher for DTW distances (**Supporting Information S2**). Among the crisp CVIs, a majority of 6 out of the 7 indices (Sil, SF, DB, DBStar, D, COP) were consistently higher for the partitional k-means clustering with SBD distances, and this method was therefore selected for empirical time-series clustering. The most stable solutions were obtained at *k* = 6 and *k* = 7, beyond which validity scores dropped sharply. Since *k* = 7 scored lower than *k* = 6 for most indices (Sil, SF, DB, DB*, D), we retained six clusters as the optimal partition for all empirical clustering (more details on CVIs and the selection of k = 6 are provided in **Supporting Information S2)**.

Unsupervised clustering of WHPT time-series yielded the lowest average within-cluster distances to centroids (d = 0.019 ± 0.001), indicating compact and well-separated clusters (**Fig. 1a**). The time-series displayed distinct temporal trajectories across the 6 clusters: 68.9% of sites exhibited an increasing trajectory (W1 to W4), while 31% of sites (W5 & W6) showed a declining trend (**Fig. 1a & Fig. 4a).** Clustering of ‘Abundance’ time-series exhibited higher within-cluster distances (d = 0.05 ± 0.008), reflecting more heterogeneous groupings **(Fig. 2a)**. Clear increasing trajectory was obtained only for A1 (∼9% of the time-series; **Fig. 4a**), however the centroid for A2 also indicated a mild, recent recovery (8.1% sites). A strong decline was observed for A6 (16% sites), and a weak recent decline for A5 (∼20%); 46.8% of time-series (A3 & A4) exhibited complex time-series patterns. Diversity time-series also exhibited high values of within-cluster distances (d = 0.06 ± 0.015; **Fig. 3a**), with strongest cohesion within the non-linear decreasing trends of D6 (d = 0.04; 24.4% sites) and D5 (d = 0.05, 15.7% sites; **Fig 4a)**, followed by the increasing trend time-series of D1 (d = 0.07; 14.8% of the sites).

**Figure 1.**
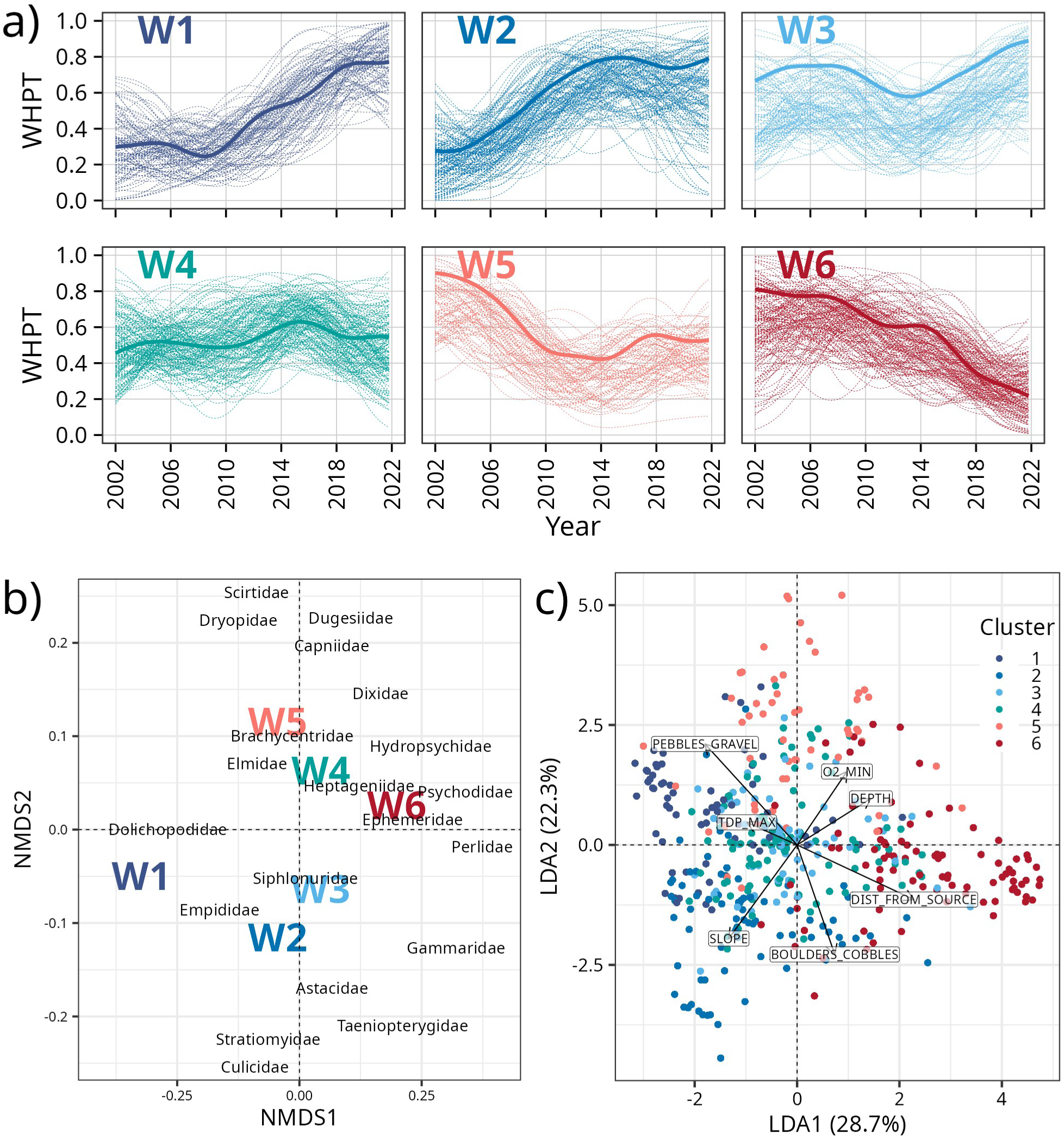
Relationships between clusters of WHPT metric time series, environmental variables, and macroinvertebrate community composition. (a) Six distinct clusters were obtained from an unsupervised clustering analysis of the WHPT metric recorded across 803 sites, between 2002-2023. Pale dotted lines represent individual time series (i.e. sites) solid thick lines represents the archetypical cluster centroid (i.e. mean temporal trend for each cluster). (b) Non-Metric Multidimensional Scaling (NMDS) plot showing variation in macroinvertebrate community composition across the six WHPT clusters (Stress = 0.033). (c) Linear Discriminant Analysis (LDA) of 803 sites based on multiple environmental variables. See **Table 1** for environmental variable definitions.

**Figure 2.**
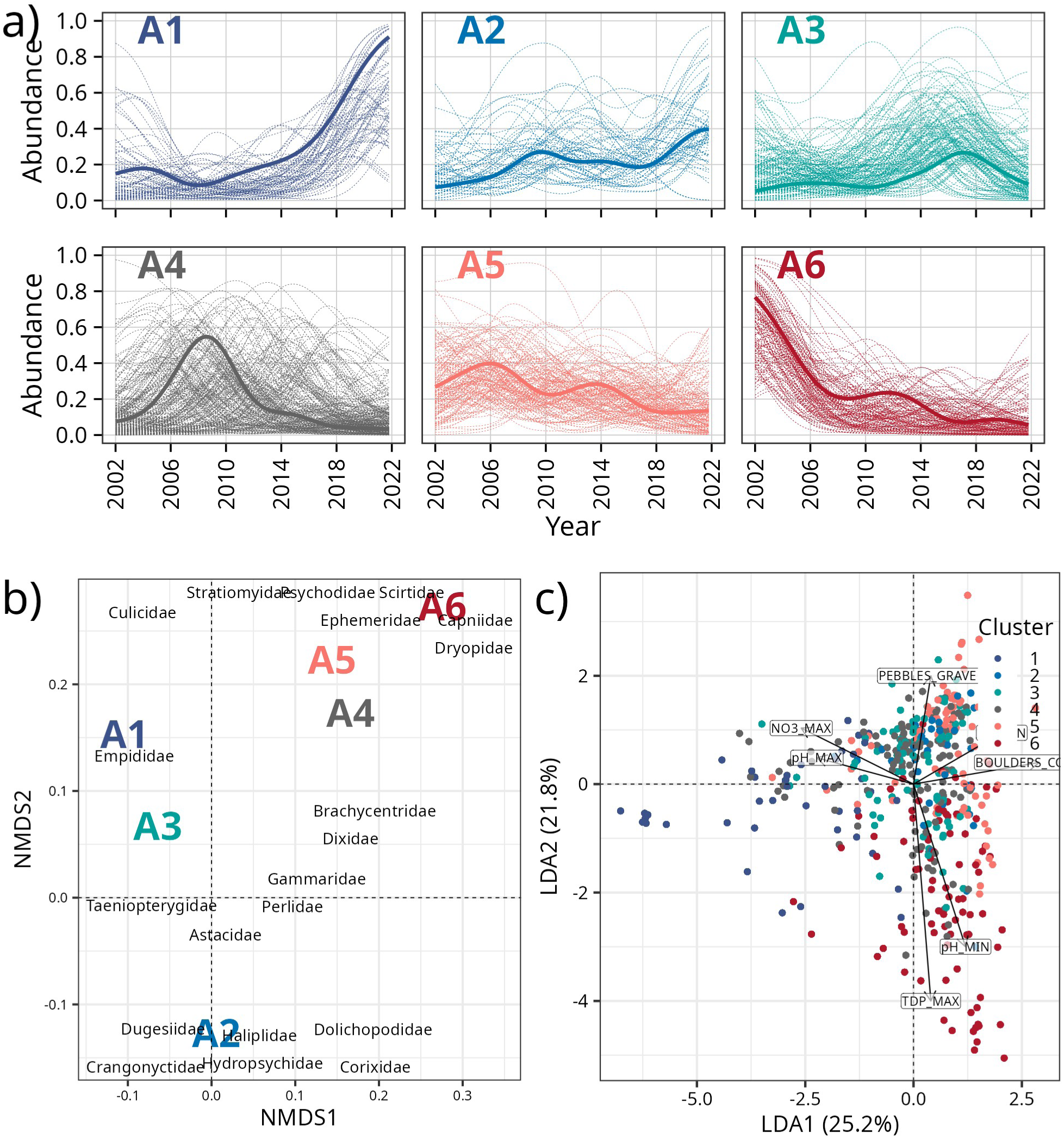
Relationships between clusters of abundance time series, environmental variables, and macroinvertebrate community composition. **(a)** Six clusters were obtained from an unsupervised clustering analysis of macroinvetebrate abundance recorded across 803 sites, between 2002-2023. Pale dotted lines represent individual time series (i.e. sites) solid thick lines represents the mean temporal trend for each cluster (i.e., the archetypical cluster centroid). **(b)** Non-Metric Multidimensional Scaling (NMDS) plot showing variation in macroinvertebrate community abundance across the six clusters (Stress = 0.041). (**c)** Linear Discriminant Analysis (LDA) of 803 sites based on multiple environmental variables (**Table 1)**

**Figure 3.**
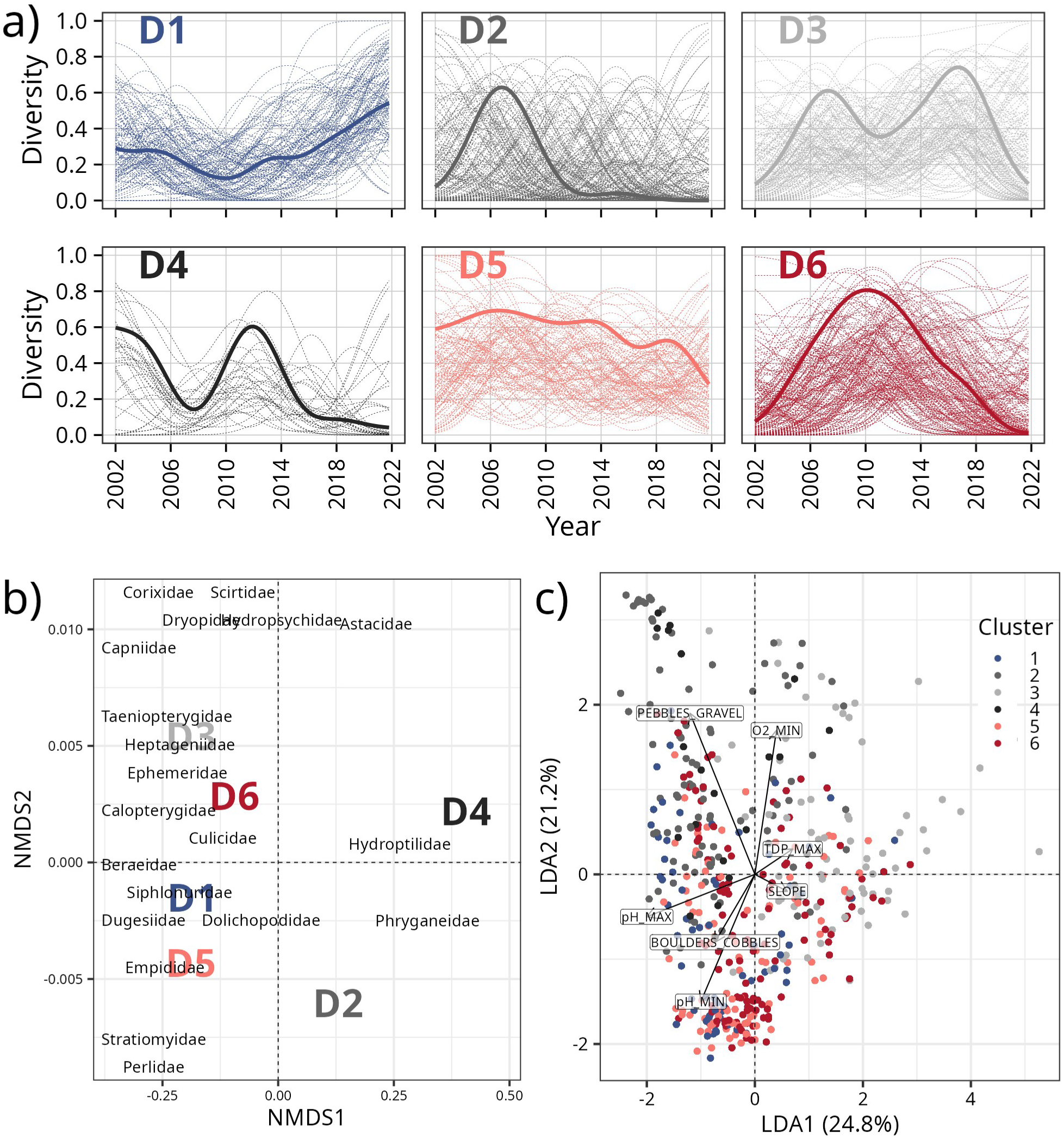
Relationships between clusters of diversity time series, environmental variables, and macroinvertebrate community composition. **(a)** Six time-series clusters were obtained from an unsupervised clustering analysis of Simpson’s Diversity metric recorded across 803 sites, between 2002-2023. Pale dotted lines represent individual time series (i.e. sites) solid thick lines represents the mean temporal trend for each cluster (i.e., the archetypical cluster centroid) **(b)** Non-Metric Multidimensional Scaling (NMDS) plot showing variation in macroinvertebrate community diversity across the six clusters (Stress = 0.02). (**c)** Linear Discriminant Analysis (LDA) of 803 sites based on multiple environmental variables (**Table 1**).

**Figure 4.**
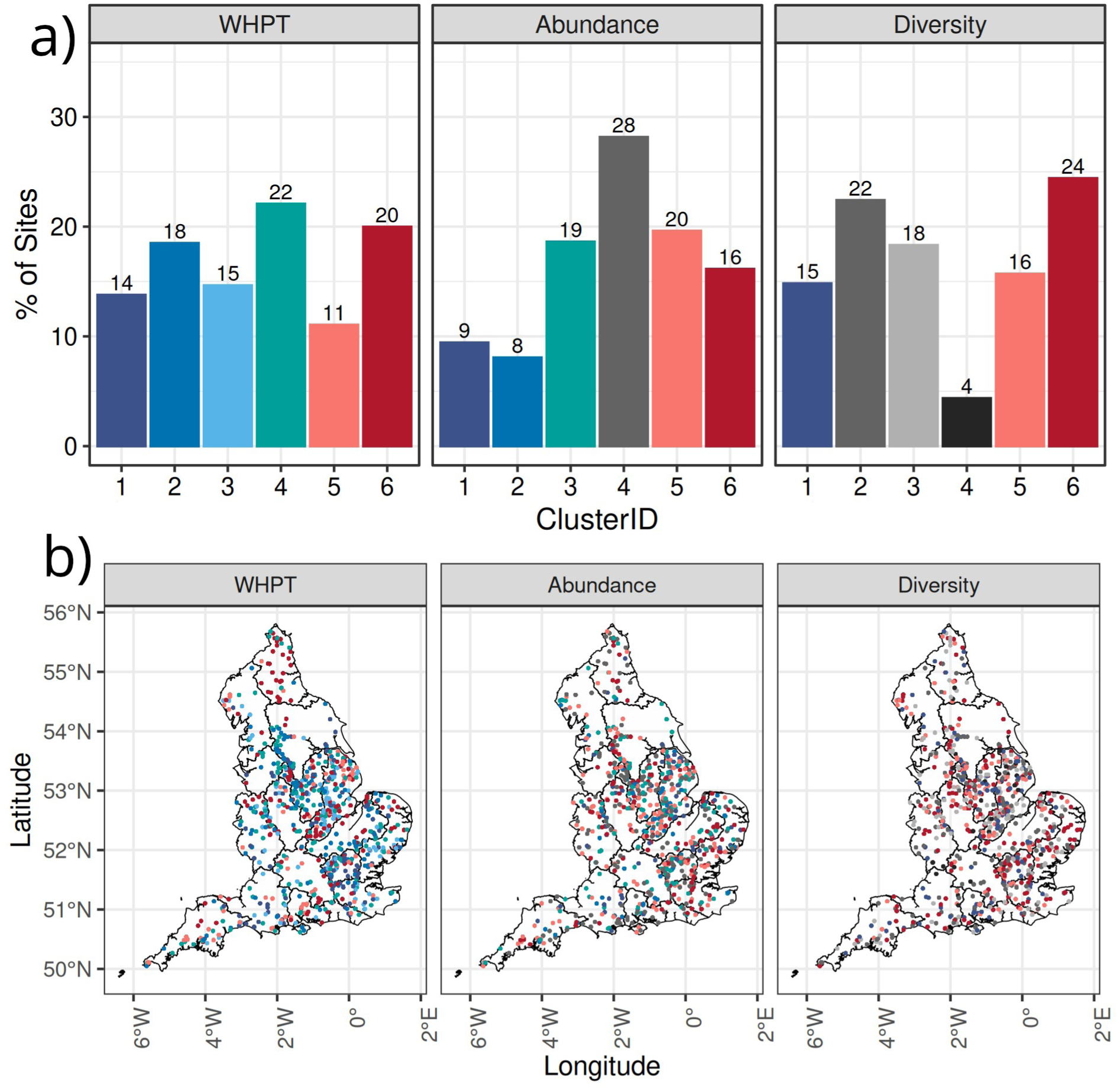
The percentages of sites per cluster, and their geographical distribution across England. **a)** Bar plots showing the percentage of sites (individual time series) assigned to different clusters. **b)** Spatial distribution of sites and clusters across England. Colors of the points are the same as cluster IDs in panel a.

All three LDA models showed strong discriminatory ability among clusters (**Table 3; Fig. S11**). The WHPT-LDA achieved the highest overall accuracy (78 - 86% across clusters; mean sensitivity ≈0.77; specificity >0.91), and the abundance- and diversity-based models performed only marginally lower (accuracy ≈70 - 81%; mean sensitivity ≈0.70–0.78; specificity mostly > 0.92; **Figs. 1c to 3c**). Cluster-specific sensitivity ranged from 0.63 to 0.80, but specificity was consistently high (>0.90 in all cases). Across all three LDAs, the total contribution of first two LD axes was relatively low. In case of WHPT clusters, LD1 and LD2 explained just half (50.9%) of total variation (LD3–LD5 = 19.2%, 16.9%, 12.9%). LD1 tracked a longitudinal position/depth–substrate gradient: increasing with DIST_FROM_SOURCE (*r* = 0.16) and DEPTH (*r* = 0.10) and decreasing with PEBBLES_GRAVEL (*r* = −0.13), while LD2 increased with PEBBLES_GRAVEL (*r* = 0.15) and with O2_MIN (*r* = 0.11), and decreased with BOULDERS_COBBLES (*r* = −0.16) and SLOPE (*r* = −0.14). In comparison, abundance and diversity clusters captured less structure (cumulative LD1 & LD2 = 47.4% and 45.8 %, respectively). LD1 contrasted coarse substrate (BOULDERS_COBBLES *r* = 0.14) against nutrient enrichment (NO3_MAX *r* = −0.12) for abundance clusters, while it contrasted O2_MIN (r = 0.12) and SLOPE (r = 0.09) with pH_MIN (r = −0.13) and DEPTH (r = - 0.07) for diversity clusters.

**Table 3.** Model Diagnostics for Linear Discriminant Analysis (LDA). Models were evaluated using three measures: sensitivity, specificity, and accuracy. Sensitivity (true positive rate) is the proportion of correctly identified positive instances out of all actual positives. Specificity (true negative rate) is the proportion of correctly identified negatives out of all actual negatives. Accuracy is the overall proportion of correctly classified instances (both positives and negatives) out of all cases. Cohen’s Kappa provides a chance-corrected measure of agreement between observed and predicted classifications, and ranges from –1 (complete disagreement) to 1 (perfect agreement). By-class statistics quantify performance for each cluster individually, and the ‘Overall statistics’ (Accuracy & Kappa) summarize performance across the entire dataset. Asterisks indicate statistical significance for the difference between observed accuracy and the no-information rate [*\*\*\* = p < 0.001; ** = p < 0.01; * = p < 0.05*]

|  |  | C1 | C2 | C3 | C4 | C5 | C6 | Accuracy | Kappa |
| --- | --- | --- | --- | --- | --- | --- | --- | --- | --- |
| WHPT | Sensitivity | 0.84 | 0.76 | 0.73 | 0.87 | 0.80 | 0.73 | 0.78 *** | 0.73 *** |
|  | Specificity | 0.76 | 0.79 | 0.78 | 0.82 | 0.78 | 0.77 |  |  |
|  | Accuracy | 0.98 | 0.97 | 0.95 | 0.97 | 0.96 | 0.91 |  |  |
| Diversity | Sensitivity | 0.72 | 0.70 | 0.80 | 0.78 | 0.75 | 0.78 | 0.75 *** | 0.69 *** |
|  | Specificity | 0.65 | 0.78 | 0.75 | 0.73 | 0.72 | 0.78 |  |  |
|  | Accuracy | 0.98 | 0.87 | 0.96 | 0.98 | 0.94 | 0.96 |  |  |
| Abundance | Sensitivity | 0.76 | 0.74 | 0.73 | 0.80 | 0.82 | 0.72 | 0.77 *** | 0.71 *** |
|  | Specificity | 0.63 | 0.77 | 0.75 | 0.81 | 0.77 | 0.76 |  |  |
|  | Accuracy | 0.99 | 0.96 | 0.95 | 0.95 | 0.93 | 0.93 |  |  |

ELSA statistics indicated poor local spatial auto-correlations for all three metrics (mean elsa >= 0.65; **Fig. 4b; also see Figs. S12-S15**). Enterograms suggested that spatial structure was strongest at neighborhood scales below 1 km for all three metrics (**Fig. S12)**.

Tests of multivariate dispersion showed no significant differences in within-cluster variability for WHPT (F=1.50, p=0.19), Abundance (F=1.13, p=0.35), or Diversity (F=1.91, p=0.09) (**Table 4; Fig. S16**). PERMANOVA indicated significant separation of communities among clusters for all three metrics (WHPT: R² = 0.34, F = 5.46, p < 0.01; Abundance: R²=0.11, F = 4.79, p < 0.05; Diversity: R² = 0.08, F = 2.42, p < 0.05), and pairwise comparisons were significant for the increasing and decreasing clusters of WHPT (notably W1,W2, & W3 versus W5 & W6; adjusted p < 0.05) and Abundance (A2 vs A5 & A6; adjusted p < 0.001). (**Fig. 1b - 3b; Table 4**; NMDS Stress < 0.041 across all three metrics).

**Table 4.** Statistics from the multivariate dispersion (betadisper) and permutational multivariate analysis of variance. (**PERMANOVA) tests of family-level taxonomic composition across the six unsupervised time-series clusters of WHPT, Abundance, and Diversity metrics.** While betadisper evaluates whether the within-cluster dispersion (variance in community composition) differs significantly among clusters, PERMANOVA tests whether centroids of multivariate community composition differ among clusters (i.e. whether overall assemblage structure is more similar within than between clusters). R² values represent the proportion of compositional variance explained by cluster identity. Significant pairwise contrasts (*adj. p-values*) identify which cluster pairs were significant and thus driving the overall differences. [*\*\*\* = p < 0.001; ** = p < 0.01; * = p < 0.05*]

| Metric | F-statistic (betadisper) | F-statistic (PERMANOVA) | $R^2$ | Significant pairwise contrasts (adjusted) |
| --- | --- | --- | --- | --- |
| WHPT | 1.50 | 5.46<sup>***</sup> | 0.34 | (C1,C2, & C3) vs (C5 & C6) (all adj.p. < 0.03) |
| Abundance | 1.13 | 4.79<sup>*</sup> | 0.11 | C2 vs (C4, C5 & C6) (adj. p < 0.04) |
| Diversity | 1.91 | 2.42<sup>*</sup> | 0.08 | NA |

Adequate convergence was obtained for all three metric-models (R̂ = 1.00 - 1.03) with ESS spanning 1,285 - 5,670 across parameters (ESS > 1000 in all cases; **Table 5**; complete posterior predictions shown in **Fig. S17 - S19**). Posterior standard deviations of random intercepts were largest at the ‘SITE_ID’ level (3.2 - 11.6 across metrics) with a small range of 95% credible intervals (RCI < 2), indicating that the largest share of unexplained heterogeneity in all three metrics arises at the local site level. The second largest posterior SDs were obtained for the cross-classified ‘Typology’ level (SD = 0.5 - 1.9; RCI ≈ 5), followed by ‘Catchment’ (SD = 0.8 – 2.4), ‘Water Body’ levels (SDs = 0.9 - 2.1; RCIs = 5 – 12), and then ‘Agency-area’ level (SD = 0.7 - 2.3; RCI = 5 – 11) (**Fig. 5**).

**Figure 5.**
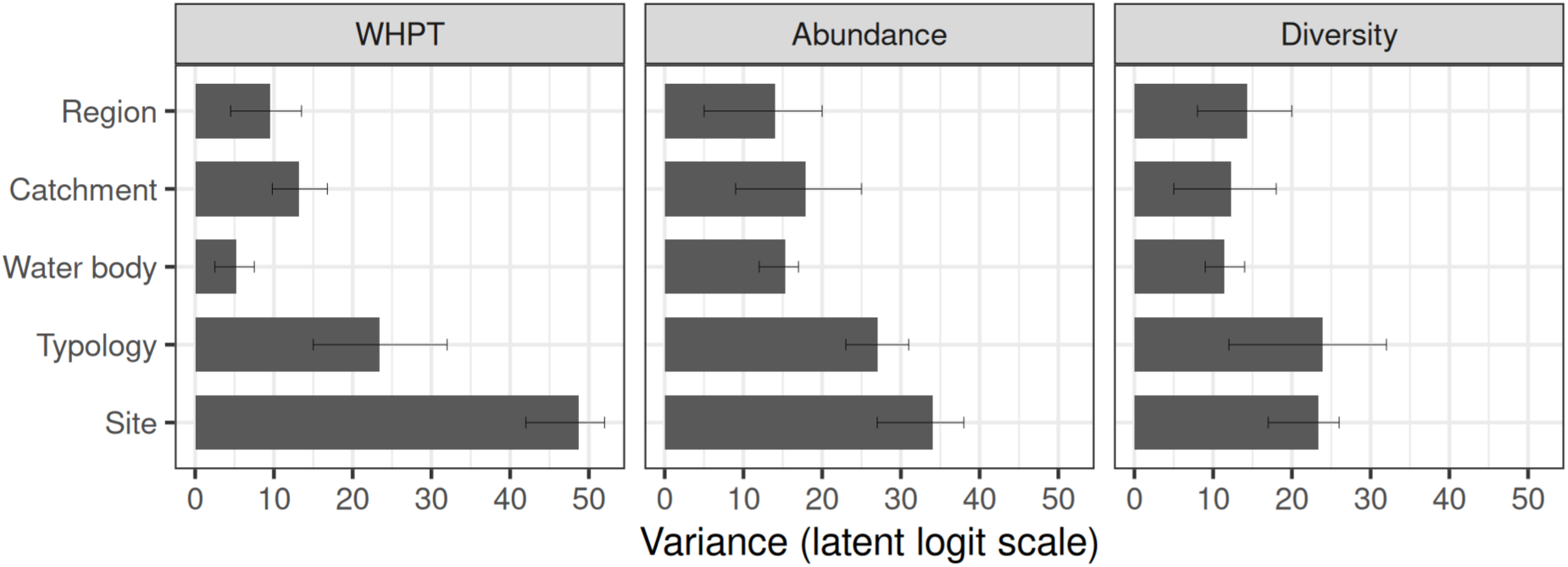
Posterior variance partitioning derived from Bayesian categorical models. Cluster variability was partitioned across hierarchical and cross-linked spatial scales (site, water body, catchment, typology, and Environment Agency region) for three metrics (WHPT, abundance, diversity). Bars show posterior means of variance components on the latent logit scale (the relative magnitude of heterogeneity in Cluster ID, attributable to each spatial level), with error bars representing 95% credible intervals.

**Table 5.** Population-level (fixed) intercepts from Bayesian categorical (multinomial-logit) models of cluster membership for time-series of WHPT, Abundance, and Diversity. Coefficients are log-odds for the six categories (C_k_) relative to the reference category, reported as posterior mean (Estimate), posterior SD (Est.Error), and 95% credible intervals (l-95% CI, u-95% CI). R̂ is the split-chain convergence diagnostic (values ≈1 indicate good mixing). Bulk-ESS and Tail-ESS are effective sample sizes for central and tail posterior summaries, respectively. For each metric, the reference category (i.e. the baseline) was set as the cluster where time-series exhibited neither a strong increasing or decreasing trend.

| | | Estimate ($\mu$) | Est.Error | l-95%<br>CI | u-95%<br>CI | $\hat{R}$ | Bulk_ESS | Tail_ESS |
| --- | --- | --- | --- | --- | --- | --- | --- | --- |
| **WHPT** | **C1** | -2.64 | 1.59 | -6.62 | -0.68 | 1.00 | 1236 | 1558 |
|  | **C2** | -3.21 | 1.88 | -7.87 | -0.95 | 1.00 | 804 | 1146 |
|  | **C3** | -2.77 | 1.83 | -7.3 | -0.62 | 1.01 | 848 | 757 |
|  | **C4** | -2.07 | 1.45 | -5.58 | -0.26 | 1.00 | 752 | 1350 |
|  | **C5** | -1.86 | 1.39 | -5.43 | -0.26 | 1.01 | 688 | 711 |
| **Abundance** | **C2** | -1.89 | 2.28 | -7.61 | 1.21 | 1.01 | 2648 | 1758 |
|  | **C3** | -1.62 | 2 | -6.49 | 0.94 | 1.00 | 1848 | 817 |
|  | **C4** | -2.25 | 1.94 | -7.03 | 0.06 | 1.01 | 1808 | 1509 |
|  | **C5** | -2.66 | 2.81 | -9.34 | 0.65 | 1.01 | 1852 | 620 |
| **Diversity** | **C2** | -1.5 | 2.2 | -7.19 | 1.29 | 1.02 | 1028 | 695 |
|  | **C3** | -1.42 | 2.27 | -7.17 | 1.36 | 1.00 | 956 | 793 |
|  | **C4** | -1.07 | 2.04 | -5.89 | 1.8 | 1.03 | 1092 | 1076 |
|  | **C5** | 0.23 | 1.87 | -4.04 | 3.1 | 1.02 | 1204 | 1297 |
|  | **C6** | -1.25 | 2.13 | -6.55 | 1.39 | 1.00 | 1308 | 829 |

## Discussion

Across more than 800 long-term monitoring sites, temporal trends were highly heterogeneous, ranging from strong increases to pronounced declines. Shape-explicit time-series analysis revealed distinct groups (clusters) of sites, representing divergent ecological pathways of recovery and decline that were strongly structured by longitudinal position and river typology. Site-level variation dominated, such that sites within the same region, catchment, or waterbody differed more from each other than regions, catchments, or waterbodies did from one another. Nonetheless, river typology accounted for substantial variation in time-series shapes, underscoring the role of variables such as catchment size, dominant geology and altitude in mediating or modulating recovery dynamics. Together, these findings demonstrate that freshwater biodiversity change in UK rivers cannot be reduced to a single national trajectory. Instead, it reflects a mosaic of asynchronous, site-specific pathways. The spatial structure in these patterns highlights the influence of network position and connectivity, conditioned by typology, on long-term ecological trajectories. We elaborate on these patterns and their mechanistic implications in the sections that follow.

### Complex and asynchronous temporal dynamics

Consistent with our first hypothesis, freshwater macroinvertebrate communities did not follow simple monotonic trajectories. Shape-based clustering revealed a wide range of temporal patterns, with approximately half of all site–metric combinations (≈51%) showing complex, non-linear dynamics. Among the metrics, WHPT exhibited the strongest and most coherent clustering structure. This suggests that pollution-sensitivity scores provide a more stable signal of ecological change as compared to abundance or diversity metrics. WHPT’s relative robustness likely reflects its logarithmic scoring system, which reduces its sensitivity to sampling and demographic stochasticity in natural populations. By contrast, abundance and diversity trajectories were noisier and more weakly structured. For these metrics, robust inference is hampered not only by demographic fluctuations, but also by uncertain historical baselines, imperfect detection, taxonomic aggregation, and artefacts associated with density dependence – challenges well recognised in the biodiversity-monitoring literature (Didham et al., 2020; Wilkes et al., 2025a; Wilkes et al., 2025b). Diversity patterns were especially diffuse, with high within-cluster dispersion and weak archetypal signatures; richness–evenness indices are inherently sensitive to rare species dynamics, shifts in dominance structure, and incomplete taxonomy (Jost, 2006; Chao et al., 2014; Wilkes et al., 2025b). Owing to this contrast, coupled with the noisier across-cluster partitions for abundance and diversity metrics, we have concentrated more on discussing the ecological underpinnings of the WHPT clusters in the remaining sections, as these provide more diagnostically interpretable ecological groupings.

Traditional slope- or effect-size–based approaches would classify all pollution-sensitivity (WHPT) time-series with early plateaus or delayed rebounds simply as “increasing” or “decreasing,” thereby obscuring the temporally varying shapes that reveal underlying ecological processes and management effects. For instance, shape-based clustering delineated two clusters (W1 and W2) that both exhibited increasing trajectories but differed markedly in the timing and form of recovery. Sites within W1 showed an early plateau until ∼2008, followed by a steep rise until ∼2018 and a subsequent levelling-off, whereas W2 displayed a steady increase until ∼2015 before reaching a plateau. Both clusters likely reflect responses to improving water quality, yet their divergent shapes indicate distinct recovery pathways. These differences may arise because (i) the mitigation of chronic stressors, and thus the onset of recovery, occurred at different times across sites, or (ii) both clusters responded differently to similar management interventions due to variations in network connectivity, baseline community composition, or both, suggesting contrasting ecological routes to recovery (Brown and Swan, 2010; Tonkin et al., 2016; Tonkin et al., 2018; Van Looy et al., 2019). For instance, our LDA analysis indicates that W1 sites are closer to their longitudinal sources along the network, as compared to W2 sites. Thus the initial plateau in W1 can indicate delayed recovery driven by dispersal limitation within a weakly connected metacommunity (Tonkin et al., 2018; Borthagaray et al., 2020). In contrast, W2 sites may have experienced faster rescue effects once conditions improved, leading to a more immediate rise without a temporal lag (Borthagaray et al., 2020). Similarly, although both clusters ultimately exhibit late plateaus, their timing differed markedly. The later levelling-off (∼2018) of the poorly connected W1 sites may result from (i) the buffering influence of a more functionally resilient baseline assemblage (Van Looy et al., 2019) and/or (ii) a deceleration of recovery due to residual or emerging stressors (Haase et al., 2023). Conversely, the earlier levelling (∼2015) of the W2 sites may reflect a more functionally homogeneous source pool, which reached saturation once the limits of ecological niches or functional redundancy were attained (Lamothe et al., 2018). Overall, these trajectories highlight far more complex temporal dynamics than the uniform plateau reported by Haase et al. (2023). If UK results are representative of wider European results as suggested by Haase et al. (2023), the apparent pan-European “halt” in biodiversity recovery likely conceals considerable local heterogeneity in both timing and ecological pathways.

A similar divergence was evident among the declining trajectories: in W5, declines halted around 2010 and even reversed slightly after ∼2014, whereas W6 showed a more monotonic decline. The partial rebound in W5 likely reflects site-specific mitigation of chronic stressors (e.g. reductions in agricultural runoff or industrial discharges), or improved connectivity to upstream source pools through morphological restoration (Van Looy et al. 2019; Whelan et al. 2022). It is most likely that multiple interacting processes underlie the deviation between W5 and W6, and disentangling their relative influence will be critical for sustaining biodiversity gains amid continuing environmental pressures (Vaughan & Ormerod 2012, 2014; Verbeck et al. 2016). As policy frameworks increasingly seek to align biodiversity monitoring with ecological restoration targets (Wilkes et al. 2025a), shape-explicit, trajectory-based analyses provide a more sensitive and therefore mechanistically grounded diagnostic lens for identifying when, where, and why recovery stalls or reverses.

### Environmental drivers and community composition across the clusters

The coexistence of complex trajectories confirms that freshwater biodiversity change is shaped by multiple, interacting factors. Although we cannot directly attribute temporal trends within individual clusters to specific environmental variables due to spatial and temporal mismatches in the availability of biological, hydrological and water-quality data, LDA on River Invertebrate Classification Tool (RICT) drivers provided correlational support for this interpretation. The separation between clusters of increasing and decreasing trends was primarily structured by a composite gradient of longitudinal position, substrate composition, water depth, and channel slope, in line with our second hypothesis (H2). The strong influence of longitudinal position suggests a possible source–sink dynamic, with recovery trends associated with upland and mid-catchment sites and declines prevailing in downstream reaches and deeper waters. Although upland recovery and spatial source–sink structuring are well established (Ormerod and Durance, 2009; Shilland et al., 2015; Stockdale et al., 2014), the explicit role of network-scale source–sink dynamics in governing temporal biodiversity trends remains largely unexplored.

Sites in the increasing WHPT clusters (W1–W3; and to a lesser extent W4) were typically shallower, steeper, and dominated by coarse substrates, characteristic of upland and mid-catchment rivers. These sites likely function as potential source populations or species pools: their coarse substrate and high aeration promote both persistence during stress and rapid recovery when improving conditions allow. Recovery in mid-reaches is likely driven by network-mediated recolonization: sites with good connectivity to adjacent source pools can benefit from rescue effects and faster recolonization (e.g. Sundermann et al., 2011; Stoll et al., 2014; Stoll et al., 2016). In contrast, the decreasing trajectories in the relatively downstream clusters (W5–W6) appear to function as sink regions: here point-source stressors (e.g. effluent discharge, nutrient-rich run-off, chemical spill) as well as longitudinal dispersal constraints (channelization, impoundments, culverting) accumulate, resulting in a failure of recovery and recolonization (Tonkin et al., 2018; Savary et al., 2025).

The segregation between W5 and W6 along the longitudinal gradient provides strong evidence that landscape connectivity and dispersal constraints are major drivers of local community trajectories. Although both clusters represent ‘sink’ conditions typical of downstream reaches, they differ starkly in their relative longitudinal positions along the river network. Here, distance-from-source is not only a geomorphological descriptor but also serves as a proxy for proximity to upstream species source pools, where sensitive taxa are more abundant and colonist supply is expected to be higher. W6 sites occur further downstream, meaning they are more isolated from potential sources of pollution-sensitive colonists (Demars et al., 2014). In addition, they are disproportionately exposed to cumulative catchment stressors and often occur within more degraded landscape matrices. This combination of restricted connectivity and strong trait–environment matching constrains recolonisation potential producing trajectories of continued decline rather than recovery or plateauing (but see Brown and Swan, 2010). In contrast, W5 sites are located closer to source pools, which likely explains their partial recovery. Their proximity enhances dispersal opportunities and exposes them to fewer accumulated stressors than the more distal W6 sites. Overall, our results suggest that in addition to water quality and hydrological drivers, long-term community trajectories could be influenced by connectivity patterns that regulate dispersal and recolonization within dendritic river networks (Carrara et al., 2012; Carrara et al., 2015; Altermatt et al., 2013; Tonkin et al., 2018; Carraro et al., 2023). These meta-community drivers of long-term biological dynamics need to be unpicked further with process-based modelling frameworks (Thompson et al., 2020).

It is important to note that the separation of increasing and decreasing trajectories along the longitudinal gradient was not absolute. Considerable overlap was evident across the environmental space. Such deviations likely arise from a range of other drivers including: natural geomorphic adjustments (Rice et al., 2001), legacy or localised stressors (e.g. episodic pollution, agricultural and urban runoff, nutrient or sediment loading; Townsend, 1989; Palmer et al., 1997; Benda et al., 2004), morphological alterations (e.g. dams, weirs, and channel realignments; see Serial Discontinuity Concept; Ward and Stanford, 1983, 1995), and/or macroscale climatic forcing (e.g. North Atlantic Oscillation (NAO); Durance and Ormerod, 2007, 2009; Larsen et al., 2024). Indeed, our analysis revealed that several oscillatory clusters (W1, W2, and W4) were positively correlated with the winter NAO index, while one cluster (W6) showed a significant negative correlation (**Supporting Information S5**), indicating that large-scale climatic oscillations can synchronise or desynchronise local community trajectories (Bradley and Ormerod, 2001). Together, these factors create a network that does not act as a uniform source–sink longitudinal gradient, but a mosaic of partially connected habitats (Pringle, 2001, 2003, 2006), which are further modulated by broad-scale climatic forces (Bradley and Ormerod, 2001; Durance and Ormerod, 2007, 2009; Larsen et al., 2024). Incorporating these localised departures, driven by legacy stressors, disrupted connectivity, and climatic variability, is essential for biodiversity monitoring frameworks that aim not only to detect change but to diagnose its underlying mechanisms across scales.

The different clusters did not fully resolve in their taxonomic composition, and the NMDS patterns deviate from Hypothesis H3, underscoring the inherent complexity of freshwater community responses along the longitudinal gradient. Both sensitive and tolerant taxa occurred within clusters linked to increasing and decreasing WHPT trajectories. This overlap also helps explain why some sites exhibited increasing total abundance but declining WHPT scores, and vice-versa (e.g. Jones et al. 2023; McKenzie et al. 2022; Powell et al. 2023; Johnson et al. 2024). These metrics capture different ecological dimensions: total abundance reflects overall productivity and numerical dominance, whereas WHPT represents the relative contribution of pollution-sensitive taxa. Hence, abundance may rise due to the proliferation of tolerant groups even as sensitive EPT taxa decline, while WHPT may increase when sensitive taxa recolonise but overall counts remain low due to habitat or seasonal constraints. Similar divergences occur between WHPT and diversity, or between abundance and diversity, as the diversity indices are primarily influenced by evenness rather than total abundance or tolerance weighting. Together, these patterns emphasise that productivity, sensitivity, and evenness respond differently to environmental change, and therefore trends derived from different metrics are not directly comparable.

### Scaling ecological variability: from local patches to river networks

Across our analyses, a consistent result was the absence of a single, coherent geographic structure in trajectory clusters. Variation in trajectories was greater among individual sites than among catchments, agency areas, or even typological classes, in line with hypothesis H4. Some previous studies noted that individual sites can differ in their temporal responses (Vaughan and Ormerod, 2012; Larsen et al., 2019; Johnson et al., 2019; Powell et al., 2023), but to our knowledge no study has explicitly quantified the relative variance across the full spatial hierarchy. While we did not detect strong spatial autocorrelation at neighbourhood scales, we did not account for flow connectivity, flow direction, or network topology in the autocorrelation calculations, although these factors are well known to affect ecological similarity in river networks (Carrara et al., 2012; Altermatt et al., 2013). Even without constructing a full network structure, our ELSA Moran or local autocorrelation/enterogram analysis revealed that sites were substantially similar in their trend shape at up to ∼1 km² spatial neighbourhood scales, beyond which cluster assignments diverged.

Second to ‘Site ID’, typology emerged as an important source of variance in temporal trends, highlighting its role as the physical and chemical template within which hydrological and ecological processes operate. To our knowledge, only one national-scale study has explicitly compared macroinvertebrate trajectories across formal river types (Powell et al., 2023). They found more positive trends across calcareous rivers (which was also the most numerous type in their dataset) than siliceous rivers. This was attributed to the greater flow, thermal, and chemical stability of groundwater-fed chalk and limestone systems relative to more surface water–dominated siliceous systems (Powell et al., 2023). Our dataset was similarly dominated by calcareous river types (System A Types II, V, and VIII; >75% of sites), with markedly fewer siliceous (∼19%) and organic (∼2%) systems. Despite this skewed distribution, typology still explained substantial variance (although, as expected, the relative class interval for ‘Typology’ (RCI ≈ 5) was larger than ‘SITE_ID’ (RCI < 2), since the contrasting types are sparsely sampled). Even a relatively small number of high-altitude, siliceous, or organic rivers followed systematically different trajectories from the dominant lowland calcareous rivers, producing strong between-typology variance. The constituent components of typology – geology (/alkalinity), altitude, flow regime – have been individually reported as influential covariates affecting invertebrate status or recovery (e.g. Vaughan and Ormerod, 2012, 2014; Monk et al., 2008; Johnson et al., 2025), yet the formal WFD/UKTAG typology classification has rarely been used as an explicit hierarchical factor. Our findings suggest that, for UK rivers, typological aggregation better reflects the ecological organisation of temporal trajectories than the more commonly used regional or catchment-level groupings, making it a more logical basis for national-scale trend assessments.

Overall, our study affirms that the most effective path for freshwater biodiversity monitoring lies not in the post-hoc aggregation of heterogeneous and opportunistic datasets, but in the development of a standardised, spatially-distributed and nested sampling network that maintains site-level resolution (preserving local ecological nuance) with frequent and consistent sampling. Thus, while long-term, site-specific monitoring remains indispensable, expanding and maintaining a sufficiently dense network would allow us to examine how changes synchronise (or fail to) across river types, typologies, catchments, and regions. In stark contrast, the ongoing contraction of monitoring capacity threatens this capability; Environment Agency funding fell from approximately £120 million in 2010 to around £40 million in 2020 (Environment Agency, 2025). Without robust, sustained investment in monitoring networks, our ability to detect, attribute and manage biodiversity change in freshwater systems will be fundamentally compromised.

## Author Contributions

**Mansi Mungee:** data curation, formal analysis, methodology, software, visualization, writing – original draft, writing – review and editing. **Lee E. Brown:** conceptualization, investigation, funding acquisition, methodology, project administration, supervision, writing – original draft, writing – review and editing. **William E. Kunin:** funding acquisition, project administration, supervision, writing – review and editing. **Christopher Hassall:** funding acquisition, investigation, project administration, supervision, writing – review and editing. **Martin Wilkes:** data, resources, writing – review and editing. **Vicky Bell:** data, resources, writing – review and editing.

## Acknowledgments

We thank Dr. Claire Carvell and Dr. D. M. Cooper (UK Centre for Ecology & Hydrology) for providing the LTLS data. M.W.’s time contribution was funded by the School of Life Sciences, University of Essex. M.M.’s time contribution was funded by the Natural Environment Research Council (NERC) Drivers and Repercussions of UK Insect Decline (DRUID) project (NE/V006916/1, NE/V006878/1) and the School of Arts and Sciences, Azim Premji University, Bhopal.

## References

1. Aghabozorgi, S., Shirkhorshidi, A.S. & Wah, T.Y. (2015) ‘Time-series clustering – a decade review.’ Information Systems, 53, 16–38. 10.1016/j.is.2015.04.007.

2. Altermatt, F., Seymour, M. & Martinez, N. (2013) ‘River network properties shape α-diversity and community similarity patterns of aquatic insect communities across major drainage basins.’ Journal of Biogeography, 40(12), 2249–2260. 10.1111/jbi.12178.

3. Anderson, M.J. (2014) ‘Permutational multivariate analysis of variance (PERMANOVA).’ Wiley StatsRef: Statistics Reference Online, 1–15. 10.1002/9781118445112.stat07841.

4. Arbelaitz, O., Gurrutxaga, I., Muguerza, J., Pérez, J.M. & Perona, I. (2013) ‘An extensive comparative study of cluster validity indices.’ Pattern Recognition, 46(1), 243–256. 10.1016/j.patcog.2012.07.021.

5. Bell, V.A., Naden, P.S., Tipping, E., Davies, H.N., Carnell, E., Davies, J.A.C. et al. (2021) ‘Long-term simulations of macronutrients (C, N and P) in UK freshwaters.’ Science of the Total Environment, 776, 145813. 10.1016/j.scitotenv.2021.145813.

6. Benda, L., Poff, N.L., Miller, D., Dunne, T., Reeves, G., Pess, G. & Pollock, M. (2004) ‘The network dynamics hypothesis: how channel networks structure riverine habitats.’ BioScience, 54(5), 413–427. 10.1641/0006-3568(2004)054[0413:TNDHHC]2.0.CO;2.

7. Bloomfield, N.J., Knerr, N. & Encinas-Viso, F. (2018) ‘A comparison of network and clustering methods to detect biogeographical regions.’ Ecography, 41(1), 1–10. 10.1111/ecog.02810.

8. Borthagaray, A.I., Teixeira-de Mello, F., Tesitore, G., Ortiz, E., Illarze, M., Pinelli, V., Urtado, L., Raftopulos, P., González-Bergonzoni, I., Abades, S. & Loureiro, M. (2020) ‘Community isolation drives lower fish biomass and species richness, but higher functional evenness, in a river metacommunity.’ Freshwater Biology, 65(12), 2081–2095. 10.1111/fwb.13603.

9. Bradley, D.C. & Ormerod, S.J. (2001) ‘Community persistence among stream invertebrates tracks the North Atlantic Oscillation.’ Journal of Animal Ecology, 70(6), 987–996. 10.1046/j.0021-8790.2001.00559.x.

10. Brown, B.L. & Swan, C.M. (2010) ‘Dendritic network structure constrains metacommunity properties in riverine ecosystems.’ Journal of Animal Ecology, 79(3), 571–580. 10.1111/j.1365-2656.2010.01668.x.

11. Bürkner, P.C. (2017) ‘brms: An R package for Bayesian multilevel models using Stan.’ Journal of Statistical Software, 80(1), 1–28. 10.18637/jss.v080.i01.

12. Bürkner, P.C. (2021) ‘Bayesian item response modeling in R with brms and Stan.’ Journal of Statistical Software, 100(5), 1–54. 10.18637/jss.v100.i05.

13. Cano-Barbacil, C., Sinclair, J.S., Welti, E.A. & Haase, P. (2025) ‘Recovery and degradation drive changes in the dispersal capacity of stream macroinvertebrate communities.’ Global Change Biology, 31(1), e70054. 10.1111/gcb.70054.

14. Carrara, F., Altermatt, F., Rodriguez-Iturbe, I. & Rinaldo, A. (2012) ‘Dendritic connectivity controls biodiversity patterns in experimental metacommunities.’ Proceedings of the National Academy of Sciences, 109(15), 5761–5766. 10.1073/pnas.1119651109.

15. Carrara, F., Giometto, A., Seymour, M., Rinaldo, A. & Altermatt, F. (2015) ‘Inferring species interactions in ecological communities: a comparison of methods at different levels of complexity.’ Methods in Ecology and Evolution, 6(8), 895–906. 10.1111/2041-210X.12382.

16. Carraro, L., Hartikainen, H., Jokela, J., Bertuzzo, E. & Rinaldo, A. (2023) ‘Modelling environmental DNA transport in rivers reveals scale-dependent biodiversity patterns.’ Scientific Reports, 13, 8885. 10.1038/s41598-023-35614-6.

17. Carrier-Belleau, C., Pascal, L., Tiegs, S.D., Nozais, C. & Archambault, P. (2023) ‘Tipping point arises earlier under a multiple-stressor scenario.’ Scientific Reports, 13(1), 16780. 10.1038/s41598-023-43738-1.

18. Chao, A., Gotelli, N.J., Hsieh, T.C., Sander, E.L., Ma, K.H., Colwell, R.K. & Ellison, A.M. (2014) ‘Rarefaction and extrapolation with Hill numbers: a framework for sampling and estimation in species diversity studies.’ Ecological Monographs, 84(1), 45–67. 10.1890/13-0133.1.

19. Chesters, R.K. (1980) Biological monitoring working party: the 1978 national testing exercise. Technical Memorandum No. 19. Department of the Environment, Water Data Unit.

20. Civan, A., Worrall, F., Jarvie, H.P., Howden, N.J. & Burt, T.P. (2018) ‘Forty-year trends in the flux and concentration of phosphorus in British rivers.’ Science of the Total Environment, 640–641, 1524–1535. 10.1016/j.scitotenv.2018.05.392.

21. Demars, B.O., 2024. European freshwater macroinvertebrate richness and abundance: alternative analyses and new findings. bioRxiv, pp.2024–08. 10.1101/2024.08.13.607735

22. Demars, B.O., Wiegleb, G., Harper, D.M., Bröring, U., Brux, H. & Herr, W. (2014) ‘Aquatic plant dynamics in lowland river networks: connectivity, management and climate change.’ Water, 6(4), 868–911. 10.3390/w6040868.

23. Didham, R.K., Basset, Y., Collins, C.M., Leather, S.R., Littlewood, N.A., Menz, M.H., Müller, J., Packer, L., Saunders, M.E., Schönrogge, K. & Stewart, A.J. (2020) ‘Interpreting insect declines: seven challenges and a way forward.’ Insect Conservation and Diversity, 13(2), 103–114. 10.1111/icad.12408.

24. Directive, W.F. (2000) ‘EU Water Framework Directive.’ EC Directive, 60.

25. Doretto, A., Piano, E. & Larson, C.E. (2020) ‘The River Continuum Concept: lessons from the past and perspectives for the future.’ Canadian Journal of Fisheries and Aquatic Sciences, 77(11), 1853–1864. 10.1139/cjfas-2020-0080.

26. Dudgeon, D., Arthington, A.H., Gessner, M.O., Kawabata, Z.-I., Knowler, D.J., Lévêque, C., Naiman, R.J., Prieur-Richard, A.-H., Soto, D., Stiassny, M.L. & Sullivan, C.A. (2006) ‘Freshwater biodiversity: importance, threats, status and conservation challenges.’ Biological Reviews, 81(2), 163–182. 10.1017/S1464793105006950.

27. Durance, I. & Ormerod, S.J. (2009) ‘Trends in water quality and discharge confound long-term warming effects on river macroinvertebrates.’ Freshwater Biology, 54(2), 388–405. 10.1111/j.1365-2427.2008.02112.x.

28. Durance, I. & Ormerod, S.J. (2007) ‘Climate change effects on upland stream macroinvertebrates over a 25-year period.’ Global Change Biology, 13(5), 942–957. 10.1111/j.1365-2486.2007.01340.x.

29. Environment Agency (2014) Freshwater macro-invertebrate analysis of riverine samples. LIT 11614

30. Environment Agency (2017) Freshwater macro-invertebrate sampling in rivers. LIT 11610

31. Environment Agency (2025) Annual report and accounts for the financial year 2024 to 2025 at www.gov.uk/official-documents

32. Fornaroli, R., White, J.C., Boggero, A. & Laini, A. (2020) ‘Spatial and temporal patterns of macroinvertebrate assemblages in the River Po Catchment (Northern Italy).’ Water, 12(9), 2452. 10.3390/w12092452.

33. Freeman, M.C., Pringle, C.M. & Jackson, C.R. (2007) ‘Hydrologic connectivity and the contribution of stream headwaters to ecological integrity at regional scales.’ JAWRA Journal of the American Water Resources Association, 43(1), 5–14. 10.1111/j.1752-1688.2007.00002.x.

34. Greenop, A., Woodcock, B.A., Outhwaite, C.L., Carvell, C., Pywell, R.F., Mancini, F., Edwards, F.K., Johnson, A.C. & Isaac, N.J.B. (2021) ‘Patterns of invertebrate functional diversity highlight the vulnerability of ecosystem services over a 45-year period.’ Current Biology, 31(20), 4627–4634. 10.1016/j.cub.2021.08.060.

35. Halsband, C. & Sørensen, L. (2020) ‘Microplastics in the aquatic environment…’ Environmental Toxicology and Chemistry, 39(6), 1119–1134. 10.1002/etc.4718.

36. Hawkes, H.A. (1998) ‘Origin and development of the biological monitoring working party score system.’ Water Research, 32(3), 964–968. 10.1016/S0043-1354(97)00275-3.

37. Hegg, J.C. & Kennedy, B.P. (2021) ‘Let’s do the time warp again…’ Ecosphere, 12(9), e03742. 10.1002/ecs2.3742.

38. Heino, J., Melo, A.S., Siqueira, T., Soininen, J., Valanko, S. & Bini, L.M. (2015) ‘Metacommunity organisation…’ Freshwater Biology, 60(5), 845–869. 10.1111/fwb.12533.

39. Hernández Martínez de la Riva, A., Harper, M., Rytwinski, T., Sahdra, A., Taylor, J.J., Bard, B., Bennett, J.R., Burton, D., Creed, I.F., Haniford, L.S. & Hanna, D.E. (2023) ‘Tipping points in freshwater ecosystems: an evidence map.’ Frontiers in Freshwater Science, 1, 1264427. 10.3389/ffwsc.2023.1264427.

40. Johnson, A.C., Jürgens, M.D., Edwards, F.K., Scarlett, P.M., Vincent, H.M. & von der Ohe, P. (2019) ‘What works?…’ Environmental Toxicology and Chemistry, 38(8), 1820–1832. 10.1002/etc.4445.

41. Johnson, A., Murray-Bligh, J., Brown, L.E., Milner, A.M. and Klaar, M.J., 2024. Assessing the use of RIVPACS-derived invertebrate taxonomic predictions for river management. bioRxiv, pp.2024–06. 10.1101/2024.06.14.599001

42. Johnson, A.C., Sadykova, D., Qu, Y., Keller, V.D., Bachiller-Jareno, N., Jurgens, M.D., Eastman, M., Edwards, F., Rizzo, C., Scarlett, P.M. & Sumpter, J.P. (2025) ‘Zinc and copper have the greatest relative importance…’ Environmental Science & Technology, 59(8), 4068–4079. 10.1021/

43. Jones, J.I., Murphy, J.F., Arnold, A., Pretty, J.L., Scarlett, P.M., Wade, A.J. & Collins, A.L. (2023) ‘What do macroinvertebrate indices measure?…’ Freshwater Biology, 68(10), 2051–2065. 10.1111/fwb.14106.

44. Jost, L. (2006) ‘Entropy and diversity.’ Oikos, 113(2), 363–375. 10.1111/j.2006.0030-1299.14714.x.

45. Kaufman, L. & Rousseeuw, P.J. (2009) Finding groups in data: an introduction to cluster analysis. Hoboken, NJ: Wiley. 10.1002/9780470316801.

46. Kim, M. & Ramakrishna, R.S. (2005) ‘New indices for cluster validity assessment.’ Pattern Recognition Letters, 26(15), 2353–2363. 10.1016/j.patrec.2005.04.007.

47. Kuhn, M. (2008) ‘Building predictive models in R using the caret package.’ Journal of Statistical Software, 28(5), 1–26. 10.18637/jss.v028.i05.

48. Kuhn, M., Wing, J., Weston, S., Williams, A., Keefer, C., Engelhardt, A., Cooper, T., Mayer, Z., Kenkel, B. & R Core Team (2020) ‘Package “caret”.’ The R Journal, 12(1), 48–56

49. Lamothe, K.A., Alofs, K.M., Jackson, D.A. & Somers, K.M. (2018) ‘Functional diversity and redundancy of freshwater fish communities…’ Diversity and Distributions, 24(11), 1612–1626. 10.1111/ddi.12787.

50. Larsen, S., Joyce, F., Vaughan, I.P., Durance, I., Walter, J.A. & Ormerod, S.J. (2024) ‘Climatic effects on synchrony and stability…’ Global Change Biology, 30(1), e17017. 10.1111/gcb.17017.

51. Larsen, S., Vaughan, I.P. & Ormerod, S.J. (2019) ‘Testing the River Continuum Concept…’ Science of the Total Environment, 655, 137–146. 10.1016/j.scitotenv.2018.11.149.

52. McKenzie, M., England, J., Foster, I. and Wilkes, M., 2022. Evaluating the performance of taxonomic and trait-based biomonitoring approaches for fine sediment in the UK. Ecological Indicators, 134, p.108502. 10.1016/j.ecolind.2021.108502

53. Medupin, C. (2020) ‘Spatial and temporal variation of benthic macroinvertebrate communities…’ Environmental Monitoring and Assessment, 192(2), 84. 10.1007/s10661-019-7982-5.

54. Mining Remediation Authority (2025) Annual report and accounts 2024 to 2025: performance report www.gov.uk/official-documents

55. Monk, W.A., Wood, P.J., Hannah, D.M. & Wilson, D.A. (2008) ‘Macroinvertebrate community response to…’ River Research and Applications, 24(7), 988–1001. 10.1002/rra.1084.

56. Moritz, S. & Bartz-Beielstein, T. (2017) ‘imputeTS: Time-series missing value imputation in R.’ R Journal, 9(1), 207–218. 10.32614/RJ-2017-009.

57. Murphy, J.F., Winterbottom, J.H., Orton, S., Simpson, G.L., Shilland, E.M. & Hildrew, A.G. (2014) ‘Evidence of recovery from acidification…’ Ecological Indicators, 37, 330–340. 10.1016/j.ecolind.2013.10.026.

58. Murray-Bligh, J., 1999. Procedures for collecting and analysing macro-invertebrate samples. Environment Agency.

59. Nadaraya, E.A. (1964) ‘On estimating regression.’ Theory of Probability and Its Applications, 9(1), 141–142. 10.1137/1109020.

60. Naimi, B., Hamm, N.A.S., Groen, T.A., Skidmore, A.K., Toxopeus, A.G. & Alibakhshi, S. (2019) ‘ELSA: Entropy-based local indicator of spatial association.’ Spatial Statistics, 29, 66–88. 10.1016/j.spasta.2018.10.004.

61. Nguyen, H.H., Peters, K., Kiesel, J., Welti, E.A., Gillmann, S.M., Lorenz, A.W., Jaehnig, S.C. & Haase, P. (2024) ‘Stream macroinvertebrate communities in restored and impacted catchments…’ Science of the Total Environment, 929, 172659. 10.1016/j.scitotenv.2024.172659.

62. Oksanen, J., Kindt, R., Legendre, P., O’Hara, B., Stevens, M.H.H., Oksanen, M.J. & Suggests, M.A.S.S. (2007) ‘The vegan package.’ Community Ecology Package, 10(631–637)

63. Ormerod, S.J. & Durance, I. (2009) ‘Restoration and recovery from acidification…’ Journal of Applied Ecology, 46(1), 164–174. 10.1111/j.1365-2664.2008.01587.x.

64. Outhwaite, C.L., Gregory, R.D., Chandler, R.E., Collen, B. & Isaac, N.J.B. (2020) ‘Complex long-term biodiversity change among invertebrates, bryophytes and lichens.’ Nature Ecology & Evolution, 4(3), 384–392. 10.1038/s41559-020-1111-z.

65. Paisley, M.F., Trigg, D.J. & Walley, W.J. (2014) ‘Revision of the BMWP score system…’ River Research and Applications, 30(7), 887–904. 10.1002/rra.2686.

66. Palmer, M.A., Ambrose, R.F. & Poff, N.L. (1997) ‘Ecological theory and community restoration ecology.’ Restoration Ecology, 5(4), 291–300. 10.1046/j.1526-100x.1997.00542.x.

67. Pavoine, S., Love, M.S. & Bonsall, M.B. (2009) ‘Hierarchical partitioning of evolutionary and ecological patterns…’ Ecology Letters, 12(9), 898–908. 10.1111/j.1461-0248.2009.01344.x.

68. Pharaoh, E., Diamond, M., Ormerod, S.J., Rutt, G. & Vaughan, I.P. (2023) ‘Evidence of biological recovery from gross pollution…’ Science of the Total Environment, 878, 163107. 10.1016/j.scitotenv.2023.163107.

69. Pilotto, F., Kühn, I., Adrian, R., Alber, R., Alignier, A., Andrews, C., Bäck, J., Barbaro, L., Beaumont, D., Beenaerts, N. et al. (2020) ‘Meta-analysis of multidecadal biodiversity trends in Europe.’ Nature Communications, 11(1), 3486. 10.1038/s41467-020-17358-8.

70. Powell, K.E., Oliver, T.H., Johns, T., González-Suárez, M., England, J. & Roy, D.B. (2023) ‘Abundance trends for river macroinvertebrates vary across taxa, trophic group and river typology.’ Global Change Biology, 29(5), 1282–1295. 10.1111/gcb.16522.

71. Pringle, C.M. (2001) ‘Hydrologic connectivity and the management of biological reserves…’ Ecological Applications, 11(4), 981–998. 10.1890/1051-0761(2001)011[0981:HCATMO]2.0.CO;2.

72. Pringle, C.M. (2003) ‘What is hydrologic connectivity and why is it ecologically important?’ Hydrological Processes, 17(13), 2685–2689. 10.1002/hyp.5145.

73. Pringle, C.M. (2006) ‘The need for a more predictive understanding of hydrologic connectivity.’ Aquatic Conservation: Marine and Freshwater Ecosystems, 16(3), 227–232. 10.1002/aqc.761.

74. Qu, Y., Keller, V., Bachiller-Jareno, N., Eastman, M., Edwards, F., Jürgens, M.D., Sumpter, J.P. & Johnson, A.C. (2023) ‘Significant improvement in freshwater invertebrate biodiversity in all types of English rivers over the past 30 years.’ Science of the Total Environment, 905, 167144. 10.1016/j.scitotenv.2023.167144.

75. R Core Team (2024) R: A Language and Environment for Statistical Computing. Vienna: R Foundation. https://www.R-project.org/.

76. Ranacher, P. & Tzavella, K. (2014) ‘How to compare movement?…’ Cartography and Geographic Information Science, 41(3), 286–307. 10.1080/15230406.2014.890071.

77. Reid, A.J., Carlson, A.K., Creed, I.F., Eliason, E.J., Gell, P.A., Johnson, P.T., Kidd, K.A., MacCormack, T.J., Olden, J.D., Ormerod, S.J. & Smol, J.P. (2019) ‘Emerging threats and persistent conservation challenges for freshwater biodiversity.’ Biological Reviews, 94(3), 849–873. 10.1111/brv.12480.

78. Revenga, C., Campbell, I., Abell, R., De Villiers, P. & Bryer, M. (2005) ‘Prospects for monitoring freshwater ecosystems towards the 2010 targets.’ Philosophical Transactions of the Royal Society B, 360(1454), 397–413. 10.1098/rstb.2004.1596.

79. Rice, S.P., Greenwood, M.T. & Joyce, C.B. (2001) ‘Tributaries, sediment sources and the longitudinal organisation of macroinvertebrate fauna along river systems.’ Canadian Journal of Fisheries and Aquatic Sciences, 58(4), 824–840. 10.1139/f01-022.

80. Schmera, D., Árva, D., Boda, P., Bódis, E., Bolgovics, Á., Borics, G., Csercsa, A., Deák, C., Krasznai, E.Á., Lukács, B.A. & Mauchart, P. (2018) ‘Does isolation influence…’ Freshwater Biology, 63(1), 74–85. 10.1111/fwb.12973.

81. Schürings, C., Globevnik, L., Lemm, J.U., Psomas, A., Snoj, L., Hering, D. & Birk, S. (2024) ‘River ecological status is shaped by agricultural land use intensity across Europe.’ Water Research, 251, 121136. 10.1016/j.watres.2022.121136.

82. Shilland, E.M., Monteith, D.T., Millidine, K. and Malcolm, I.A., 2014. UK Upland Waters Monitoring Network (UKUWMN)-Scottish Sites. Annual Summary Progress Report.

83. Sinclair, J.S., Welti, E.A., Altermatt, F., Álvarez-Cabria, M., Aroviita, J., Baker, N.J., Barešová, L., Barquín, J., Bonacina, L., Bonada, N. et al. (2024) ‘Multi-decadal improvements in the ecological quality of European rivers are not consistently reflected in biodiversity metrics.’ Nature Ecology & Evolution, 8(3), 430–441. 10.1038/s41559-023-02305-4.

84. Stockdale, A., Tipping, E., Fjellheim, A., Garmo, Ø.A., Hildrew, A.G., Lofts, S., Monteith, D.T., Ormerod, S.J. & Shilland, E.M. (2014) ‘Recovery of macroinvertebrate species richness in acidified upland waters assessed with a field toxicity model.’ Ecological Indicators, 37(B), 341–350. 10.1016/j.ecolind.2013.10.026.

85. Stoll, S., Breyer, P., Tonkin, J.D., Früh, D. & Haase, P. (2016) ‘Scale-dependent effects of river habitat quality on benthic invertebrate communities…’ Science of the Total Environment, 553, 495–503. 10.1016/j.scitotenv.2016.02.126.

86. Stoll, S., Kail, J., Lorenz, A.W., Sundermann, A. & Haase, P. (2014) ‘The importance of the regional species pool…’ PLOS ONE, 9(1), e84741. 10.1371/journal.pone.0084741.

87. Sundermann, A., Stoll, S. & Haase, P. (2011) ‘River restoration success depends on the species pool of the immediate surroundings.’ Ecological Applications, 21(6), 1962–1971. 10.1890/10-0607.1.

88. Swan, C.M. & Brown, B.L. (2017) ‘Metacommunity theory meets restoration: isolation may mediate how ecological communities respond to stream restoration.’ Ecological Applications, 27(7), 2209–2219. 10.1002/eap.1602.

89. Thomas, C., Voulgarakis, A., Lim, G., Haigh, J. & Nowack, P. (2021) ‘An unsupervised learning approach to identifying blocking events…’ Weather and Climate Dynamics Discussions, 1–34. 10.5194/wcd-2021-17.

90. Tickner, D., Opperman, J.J., Abell, R., Acreman, M., Arthington, A.H., Bunn, S.E., Cooke, S.J., Dalton, J., Darwall, W., Edwards, G. & Harrison, I. (2020) ‘Bending the curve of global freshwater biodiversity loss: an emergency recovery plan.’ BioScience, 70(4), 330–342. 10.1093/biosci/biaa002.

91. Tonkin, J.D., Altermatt, F., Finn, D.S., Heino, J., Olden, J.D., Pauls, S.U. & Lytle, D.A. (2018) ‘The role of dispersal in river network metacommunities…’ Freshwater Biology, 63(1), 141–163. 10.1111/fwb.13037.

92. Tonkin, J.D., Stoll, S., Jähnig, S.C. & Haase, P. (2016) ‘Contrasting metacommunity structure and beta diversity in an aquatic-floodplain system.’ Oikos, 125(5), 686–697. 10.1111/oik.02685.

93. Townsend, C.R. (1989) ‘The patch dynamics concept of stream community ecology.’ Journal of the North American Benthological Society, 8(1), 36–50. 10.2307/1467400.

94. Van Klink, R., Bowler, D.E., Gongalsky, K.B., Swengel, A.B., Gentile, A. & Chase, J.M. (2020) ‘Meta-analysis reveals declines in terrestrial but increases in freshwater insect abundances.’ Science, 368(6489), 417–420. 10.1126/science.aax9931.

95. Van Looy, K., Tonkin, J.D., Floury, M., Leigh, C., Soininen, J., Larsen, S., Heino, J., LeRoy Poff, N., Delong, M., Jähnig, S.C. & Datry, T. (2019) ‘The three Rs of river ecosystem resilience: Resources, recruitment, and refugia.’ River Research and Applications, 35(2), 107–120. 10.1002/rra.3396.

96. Vannote, R.L., Minshall, G.W., Cummins, K.W., Sedell, J.R. & Cushing, C.E. (1980) ‘The river continuum concept.’ Canadian Journal of Fisheries and Aquatic Sciences, 37(1), 130–137. 10.1139/f80-017.

97. Vaughan, I.P. & Gotelli, N.J. (2019) ‘Water quality improvements offset the climatic debt for stream macroinvertebrates over twenty years.’ Nature Communications, 10(1), 1956. 10.1038/s41467-019-09997-1.

98. Vaughan, I.P. & Ormerod, S.J. (2012) ‘Large-scale, long-term trends in British river macroinvertebrates.’ Global Change Biology, 18(7), 2184–2194. 10.1111/j.1365-2486.2012.02662.x.

99. Vaughan, I.P. & Ormerod, S.J. (2014) ‘Linking interdecadal changes in British river ecosystems to water quality and climate dynamics.’ Global Change Biology, 20(9), 2725–2740. 10.1111/gcb.12520.

100. Verberk, W.C., Durance, I., Vaughan, I.P. & Ormerod, S.J. (2016) ‘Field and laboratory studies reveal interacting effects of stream oxygenation and warming on aquatic ectotherms.’ Global Change Biology, 22(5), 1769–1778. 10.1111/gcb.13178.

101. Vercelloni, J., Caley, M.J., Kayal, M., Low-Choy, S. & Mengersen, K. (2014) ‘Understanding uncertainties in non-linear population trajectories…’ PLOS ONE, 9(11), e110968. 10.1371/journal.pone.0110968.

102. Wagner, S., Hüffer, T., Klöckner, P., Wehrhahn, M., Hofmann, T. & Reemtsma, T. (2018) ‘Tire wear particles in the aquatic environment…’ Water Research, 139, 83–100. 10.1016/j.watres.2018.03.051.

103. Fontaine, T.D. and Bartell, S.M. eds., 1983. *Dynamics of lotic ecosystems* (pp. 494-pp). Michigan: Ann Arbor Science.

104. Ward, J.V. & Stanford, J.A. (1995) ‘Ecological connectivity in alluvial river ecosystems and its disruption by flow regulation.’ Regulated Rivers: Research & Management, 11(1), 105–119. 10.1002/rrr.3450110109.

105. Watson, G.S. (1964) ‘Smooth regression analysis.’ Sankhyā: The Indian Journal of Statistics, Series A, 26(4), 359–372. https://www.jstor.org/stable/25049340.

106. Wei, B., Xie, Y., Wang, X., Jiao, J., He, S., Bie, Q. & Duan, H. (2020) ‘Land cover mapping based on time-series MODIS-NDVI using a dynamic time warping approach…’ Land Degradation & Development, 31(8), 1050–1068. 10.1002/ldr.3510.

107. Whelan, M.J., Linstead, C., Worrall, F., Ormerod, S.J., Durance, I., Johnson, A.C., Johnson, D., Owen, M., Wiik, E., Howden, N.J. & Burt, T.P. (2022) ‘Is water quality in British rivers “better than at any time since the end of the Industrial Revolution”?’ Science of the Total Environment, 843, 157014. 10.1016/j.scitotenv.2022.157014.

108. Wilkes, M.A., Edwards, F., Jones, J.I., Murphy, J.F., England, J., Friberg, N., Hering, D., Poff, N.L., Usseglio-Polatera, P., Verberk, W.C. & Webb, J. (2020) ‘Trait-based ecology at large scales…’ Global Change Biology, 26(12), 7255–7267. 10.1111/gcb.15304.

109. Wilkes, M.A., Mckenzie, M., Johnson, A., Hassall, C., Kelly, M., Willby, N. & Brown, L.E. (2025b) ‘Revealing hidden sources of uncertainty in biodiversity trend assessments.’ Ecography, 2025(5), e07441. 10.1111/ecog.07441

110. Wilkes, M.A., Mungee, M., Naura, M., Bell, V.A. & Brown, L.E. (2025a) ‘Predicting nature recovery for river restoration planning and ecological assessment: a case study from England, 1991–2042.’ River Research and Applications, 41(1), 68–81. 10.1002/rra.4282

111. Worrall, F., Howden, N.J.K., Burt, T.P. and Jarvie, H.P., 2025. Changes in chlorophyll-a in English rivers over the last 49 years. Journal of Hydrology, p.134394. 10.1016/j.jhydrol.2025.134394

112. WWF (2024) Living Planet Report 2024 – A System in Peril. Gland, Switzerland: WWF. ISBN 978-2-88085-319-8.

