## Supplementary File for "Rising, falling, stalling: Complex biodiversity trajectories in UK rivers"

### S1. Time Series: Simulations & Clustering

#### *ARIMA modelling and simulating time-series data*

Autoregressive integrated moving average (ARIMA) model consists of three elements: autoregressive model (denoted as *p*), model integration (*d*), and moving average model (*q*). For each time-series, we identified the optimal autoregressive and moving average parameters (*p* and *q*) by iterating through multiple possible configurations. We iterated over possible values of each, *p* and *q*, from 0 to 5 (steps of 1), excluding the (0, 0) combination, to explore different models. For each combination of *p* and *q*, the model was fitted to the time-series using the R function *arima()* from the *forecast* package (v8.23.0; Hyndman et al. 2020). We used the Box-Cox transformations by setting the *lambda* parameter to 0, because no metric values could be <0.We identified the model that best fit the time-series data based on the smallest AIC value for different (*p, q*) configurations. By repeating the process for each time-series, and each metric, we obtained a distribution of *p* and *q* parameters that best describe the parameter space of our empirical dataset.

For each raw time-series, we also selected the ‘next best’ ARIMA model i.e., with the second lowest AIC values. Systematic noise was added to each optimal (*p, q*) combination using the R function *runif()* with *999* uniform values, setting the minimum and maximum *(p, q)* from the top two models, respectively. Using these values, we simulated 999 time-series of 100 time-steps each to generate a larger dataset with similar statistical properties as our empirical data. Simulations were performed using the function *simulate().* This resulted in a total of 999 randomizations x 3 metrics = 2,997 time-series of 100 time-steps each, available for testing different clustering methods within the plausible parameter space of our empirical datasets.

#### *Unsupervised time-series clustering*

Given a dataset of n time-series data *D = {F1, F2, ......, Fn}*, the process of unsupervised partitioning of *D* into *C = {C1, C2,...., Ck}*, i.e. *k* clusters, in such a way that similar time-series are grouped together based on some measure of similarity, is called time-series clustering (Aghabozorgi et al. 2015). The distance between two time-series (X and Y) is defined across all time points: dist(X,Y) = the sum of the distance between individual points *x* and *y*:

$$dist(X,Y)=\sum_{t=1}^{T}\left(dist(x_t,y_t)\right)$$

We used three different unsupervised clustering methods (hierarchical, partitioning k-means, and fuzzy c-means) and three different distance measures – Euclidean, Dynamic Time Warping (DTW), and shape-based distance (SBD) – resulting in a total of 9 different approaches (**Table S1**).

1. **Euclidean distances** are only defined for series of equal length and are sensitive to scale and time shifts (Aghabozorgi et al. 2015). They reflect similarity in time by performing a

strict one-to-one mapping between the data instances of the time series under comparison, i.e. the calculation of the Euclidean distance assumes that the i[th] point in one sequence is aligned with the i[th] point in the other

$$d_{Euc}(X,Y)=\sum_{t=1}^{T}(x_t-y_t)^2$$

2. **Dynamic Time Warping (DTW) distance** reflects similarity in "shape" by performing a one-to-many mapping, hence allowing time shifting and matching similar shapes even if they have a time-phase difference (**Fig. S3**; Aggarwal & Reddy 2014). Ecological communities exposed to the same perturbation may be broadly resilient, yet the strength and timing of their responses often differ with community composition and local context. Consequently, similar directional changes in abundance, diversity, or WHPT can appear at slightly different times across sites. To accommodate these plausible temporal lags, DTW can be a useful distance measure which assesses similarity in the overall shape of trajectories by permitting small phase shifts (**Fig. S3**). Because the exact timing of change can still be ecologically informative, we constrained the alignment with a Sakoe–Chiba band of two years (Sakoe & Chiba, 2003), limiting matches to observations within ±2 years. This ensures that distances are minimized between data points with a temporal difference of less than two years, and penalizes larger temporal shifts. (**Fig. S3**).

While there is no simple closed-form expression for DTW, it can be defined as the minimum accumulated alignment cost over all admissible warping paths, and is computed via dynamic programming. A standard formulation (length *t*, local cost $\delta(x_i,y_j)$ can be written as:

$$D_{DTW}(X,Y)=min_{\prod}\sum_{(i,j)\in\prod}^{\square}\delta(x_i,y_i)$$

where a warping path π is a sequence of index pairs from (1,1) to (t,t) satisfying boundary, monotonicity, and step constraints. More information of DTW distance can be found in Rakthanmanon (2012).

3. **Shape Based Distance (SBD)** compares the shape rather than level or scale by first z-normalizing each series and then taking the maximum normalized cross-correlation over all circular shifts (Paparrizos & Gravano, 2015). Lower SBD indicates more similar shapes. Because z-normalization removes mean and scale, SBD is well-suited when ecological interest lies in the pattern of change (rise/fall, timing) rather than absolute magnitudes; however, it is less appropriate if differences in amplitude are themselves ecologically meaningful or if strong outliers dominate the correlation. Let $\tilde{X}_t = (X_t - \tilde{X})/s_X$ and $\tilde{Y}_t = (Y_t - \tilde{Y})/s_Y$. The normalized cross-correlation at shift ω is

$$NCC_{\omega}(\tilde{X},\tilde{Y})=\frac{\sum_{t=1}^{T}\tilde{X}\tilde{Y}_{(t-\omega(modT))}}{\square}$$ and,

$$D_{SBD}(X, Y) = 1 - \max_{\omega \in 0, \ldots, T-1} NCC_{\omega}(\tilde{X}, \tilde{Y})$$

4. **Hierarchical clustering:** makes a hierarchy of clusters using agglomerative or divisive algorithms. Agglomerative algorithm considers each item as a cluster, and then gradually merges the clusters (bottom-up). In contrast, divisive algorithm starts with all objects as a single cluster and then splits the cluster to reach the clusters with one object (top-down) – ultimately, nested hierarchy of similar groups is generated based on a pair-wise distance matrix of time-series. In contrast to most algorithms, hierarchy clustering does not require the number of clusters as an initial parameter which is a well-known and outstanding feature of this algorithm. Moreover, using hierarchical clustering it is possible to cluster unequal time-series if an appropriate elastic distance measure (e.g. DTW) is used to compute the distances.
5. **Partitioning k-means clustering:** minimizes the total distance between all objects in a cluster from their cluster center (prototype). Prototype in k-Means process is defined as mean vector of time-series in a cluster. In this approach, the number of clusters, *k*, has to be pre-assigned, which is not available or feasible to be determined for many applications and diagnostic checks (e.g., the use of Cluster Validity Indices) for determining the number of clusters is not straightforward and often subjective.
6. **Fuzzy c-means clustering:** k-Means algorithms make clusters which are constructed in ‘hard’ or ‘crispy’ manner and it means that an object is either a member of a cluster or not. On the other hand, the FCM (Fuzzy c-Means) algorithm builds ‘soft’ clusters. In fuzzy clustering, an object has a degree of membership in each cluster. FCM attempts to find the most characteristic point in each cluster, which can be considered as the “centroid” of the cluster and, then, the grade of membership for each object in the clusters. Such aim is achieved by minimizing the objective function.

Before clustering, we performed a Nadaraya–Watson kernel regression smoothing of the raw time series to reduce the year-to-year noise and highlight the overall trend function (Nadaraya 1964; Watson 1964). To estimate the optimal bandwidths for smoothing (**Fig. S4**), generalized cross validation (GCV) scores were calculated by computing the density using the normal distribution implemented via the R function *dnorm()*. The GCV scores were normalized, and the *optimize()* function was used to find the bandwidth that minimizes the GCV score within a specified interval (**Fig. S5 & S6**).

All clustering was performed using the function *tsclust()* from package dtwclust (v5.5.12, Sarda-Espinosa 2017, 2018).

### S2. Time Series: Cluster Validity Indices

Cluster Validity Indices (CVI) were used to evaluate both the clustering efficiency across methods as well as the optimal number of clusters (*k*) for each method. A suite of indices is recommended to accommodate clustering across extensive time series configurations (Arbelaitz et al. 2012), and therefore we used a set of seven crisp (Sil, SF, CH, DB, DB*, D, and COP) and five fuzzy (MPC, K, T, SC, and PBMF) partitioning validity indices (see **Table S2** for more details and definitions). Most of these indices estimate the cluster cohesion (within or intra-variance) and the cluster separation (between or inter- variance) and combine them to compute a single ‘quality’ measure (Arbelaitz et al. 2012; Kim & Ramakrishna 2005). The quality scores are compared across methods to identify the optimal approach. Similarly, within each method the scores are compared across a range of *k* to identify the most efficient resolution: *k* is chosen such that adding another cluster doesn’t improve the quality score by a large fraction (Kaufman & Rousseeuw, 1990).

In the simulated dataset, fuzzy cluster validity indices (CVIs) showed heterogeneous responses across distance measures. For SBD, the highest fuzzy indices were observed for MPC (0.55 ± 0.38), with intermediate values for SC (0.49 ± 0.28) and PBMF (0.51 ± 0.24), while T (0.22 ± 0.32) and K (0.25 ± 0.40) were comparatively lower. In contrast, DTW achieved the highest values for T (0.73 ± 0.32), PBMF (0.57 ± 0.31), and K (0.47 ± 0.34), with moderate SC (0.52 ± 0.33) but lower MPC (0.33 ± 0.36). Euclidean distances showed intermediate performance, with SC (0.57 ± 0.33) and PBMF (0.54 ± 0.30) relatively strong, but inconsistent results for T (0.60 ± 0.45), MPC (0.44 ± 0.39), and K (0.13 ± 0.26) (**Fig. S7; Table S3**).

The crisp CVIs provided clearer evidence in support of SBD with partitional clustering. Averaged across *k*, partitional k-means with SBD achieved the highest Silhouette (0.56), SF (0.44), and Dunn (0.18), along with the lowest Davies–Bouldin (0.34) and DB* (0.27), compared to both hierarchical and Euclidean/DTW distances (**Fig. S8; Table S3**). COP values were also lower (0.27), indicating better within-cluster compactness. By contrast, hierarchical clustering showed weaker discrimination (Sil = 0.23–0.40; D = 0.09–0.17), and DTW- or Euclidean-based partitional clustering produced lower scores across most indices (Sil = 0.19–0.25; SF ≤ 0.13; D ≤ 0.15).

When varying the number of clusters, both *k* = 6 and *k* = 7 yielded relatively high scores across multiple indices. However, solutions with *k* > 7 showed sharp declines, including reduced Silhouette (<0.88), near-zero Dunn, and rising Davies–Bouldin values, consistent with over-partitioning (**Fig. S9; Table S3)**. Between the two optimal solutions, *k* = 6 consistently outperformed *k* = 7 on most crisp indices (Sil, SF, DB, DB*, Dunn), and was therefore selected as the optimal number of clusters for all subsequent analyses.

All biodiversity metrics were range normalized between 0 and 1 before clustering, and raw empirical time series were also smoothed using the Nadaraya–Watson kernel regression as described above (**Fig. S6**). To account for gaps in time series (missing data), we interpolated the biodiversity metrics using the linear interpolation implemented with the function *na_interpolation()*

from R package imputeTS (Moritz et al. 2017). As noted previously, we filtered data to ensure gaps for more than 3 consecutive years are not included.

### S3. Environmental Variables

To characterise local environmental conditions at each monitoring site, we assembled a suite of hydrological, water chemistry, and morphological predictors. River flow (Flow Volume; $m^3s^{-1}$) and Catchment area (CatchmentArea; on a 1km x 1km grid) were extracted from the G2G (Grid-to-Grid; Bell et al. 2018) national flow dataset. The G2G is a national-scale hydrological model for Great Britain that runs on a 1km x 1km grid (aligned with the GB national grid), at a 15-minute time-step, and is parametrised using digital datasets (e.g. soil types, land-cover; Bell et al. 2018). Annual means were computed, and aggegrated across the time-scale as mean flow across the full temporal-resolution. Thus, our LDA models assume that site to site variation in the temporal means aligns with site-to-site variation in intra-annual (i.e., seasonal) means.

We used the *Long Term Large Scale* (LTLS) model outputs from 2002–2022 (Bell et al. 2021) to estimate dissolved organic carbon (DOC), total dissolved phosphorus (TDP), nitrate ($NO_3N$), dissolved oxygen (O2), pH, and water temperature. For each of these variables, monthly rasters covering 2002–2022 were intersected with site coordinates, aggregated annually, and then summarised into long-term mean, minimum, and maximum values. Thus, each of these variables are represented by three predictors capturing central tendency and extremes at the same temporal scale as the empirical time-series (DOC_MEAN, DOC_MIN, DOC_MAX; TDP_MEAN, TDP_MIN, TDP_MAX; pH_MEAN, pH_MIN, pH_MAX; NO3_MEAN, NO3_MIN, NO3_MAX; O2_MEAN, O2_MIN, O2_MAX; and WT_MEAN, WT_MIN, WT_MAX).

For river morphology, we used Altitude, slope, distance from source, average channel width and depth, and substrate composition (percentage cover of boulders, cobbles, pebbles, and gravel) (Environmental Agency, 2003).

All variables were mean-centred and scaled to unit variance prior to analysis. Pearson's correlation coefficients were computed, which revealed several strong associations among many of the variables ($R^2$ ranging from -0.95 to +0.98; **Fig. S10**). To address this, we used variance inflation factor (VIF) analysis, which quantifies the degree of multicollinearity by measuring how much the variance of a regression coefficient is inflated due to correlations with other predictors. A VIF of 1 indicates no collinearity, while values above 5-7 are generally considered problematic, indicating severe collinearity with other predictors. We removed all variables which exceeded a threshold of 5, and further removed a few redundant and high $R^2$ variables (discharge & channel depth; Alkalinity from Environmental Agency, 2003 & pH_MEAN from LTLS dataset; O2_MEAN & O2_MAX) to improve the LDA model performance and interpretability of results. The final set of predictors retained included altitude, slope, distance from source, channel depth, boulders and cobbles, pebbles and gravel, maximum TDP, minimum and maximum pH, maximum nitrate, and minimum dissolved oxygen. These variables are expected to represent maximally independent axes of variation in hydrology, morphology, and chemistry in our dataset

Across the network of monitoring sites, these predictors spanned wide environmental gradients. Altitude ranged from just 1 m above sea level to 374 m, with a mean of 67.6 (± 64.4), while slope ranged from nearly flat (0.08°) to steep channels at 17.3° (mean 1.7 ± 2.3). Flow volume varied markedly from 0 to 87.1 $m^3/s$, averaging 3.4 ± 10.3, reflecting the inclusion of both headwater streams and larger rivers. Nutrient concentrations also varied widely: total dissolved phosphorus ranged from 0 to 0.97 mg/L (mean 0.32 ± 0.19), and nitrate from 0 to 26.3 mg/L (mean 5.8 ± 4.8). Water pH spanned from acidic (3.9) to strongly alkaline (9.0), with an average near 8.4 ± 0.9.

**S4. Classification of Sites based on River Typology**

We classified each monitoring site according to a formal river-type framework, in order to include Typology as a cross-classified grouping factor in the hierarchical Bayesian Variance Decomposition Analysis. In the United Kingdom, two typology systems can be used to classify rivers under the EU Water Framework Directive (WFD, Directive 2000): System A and System B (**Fig. S2**).

System A classifies river water bodies using fixed categorical descriptors of altitude band (< 200 m, 200–800 m, > 800 m), catchment size band (10–100 $km^2$, 100–1000 $km^2$, > 1000 $km^2$) and dominant geology (calcareous, siliceous, organic). This yields 18 discrete “GB Type” codes (**Table 2**). We adopted System A for this study because it is (i) the standard framework used by the UK Technical Advisory Group and the Environment Agency for WFD compliance, (ii) directly comparable across EU Member States owing to its fixed categorical boundaries, and (iii) provides finer classes to define ‘local’ reference conditions (Davy-Bowker et al. 2006).

As a secondary classification, we also present System B typology to facilitate direct comparison with the only other UK study that explicitly incorporated typology into models of temporal trends (Powell et al. 2023). System B simplifies the System A framework into six broad classes based on geology (calcareous, siliceous, organic) and altitude (Low < 200 m; High ≥ 200 m). This reduced scheme sacrifices some of the spatial resolution of System A but can facilitate interpretability and comparability among studies (**Fig. S2).**

For classification, altitude for each site was obtained from the Environment Agency (2023) digital elevation data, catchment area was calculated at 1 $km^2$ resolution using the grid-to-grid national hydrological dataset (MaRIUS-G2G-WAH2; Bell et al. 2018), and dominant geology was extracted from the WFD River, Canal and SWT Waterbody Classifications (2019; data.catchmentbasedapproach.org). Sites were assigned to one of the 18 “GB Type” categories defined under the UK-specific implementation of the Water Framework Directive (Directive

2000/60/EC) typology (System A; UKTAG 2003; Davy-Bowker et al. 2006). Each site retained its System A code and was assigned to a corresponding System B class based on altitude (< 200 m vs ≥ 200 m) and geology. Catchment size, included in System A, is not used in System B.

Across the 18 System A typologies, site representation ranged from 0 to 373. The most common types were Lowland, small, calcareous streams (Type II; n = 373) and Lowland, medium, calcareous rivers (Type V; n = 228), followed by Lowland, small, siliceous streams (Type I; n = 84). Several high-altitude and large-catchment categories were sparsely represented or absent (e.g., Types XV–XVIII). When aggregated under System B, the majority of sites fell within Calcareous Low (II) and Siliceous Low (VI).

**S5. Correlation between cluster trajectories and North Atlantic Oscillation (NAO) indices**

To examine whether inter-annual climatic oscillations influenced the observed temporal trajectories of macroinvertebrate communities, we tested for correlations between cluster-level time-series and the winter phase of the North Atlantic Oscillation (NAO). This analysis follows the approach of Bradley & Ormerod (2001) and Durance & Ormerod (2007, 2009), who demonstrated that NAO variability can synchronize biological responses in some UK rivers (also see Larsen et al. 2023). Cluster centroids were obtained from the six-cluster partitional, shape-based time-series analysis. Each centroid represents the smoothed (100 time-steps) composite trajectory of sites grouped by similarity in temporal community trends between 2002 and 2023. We used the Hurrell North Atlantic Oscillation (NAO) Station-Based Index (DJFM), provided by the Climate Analysis Section, NCAR, Boulder, USA (Hurrell 2003; updated 2023-03-15). The index represents the normalized sea-level pressure difference between Lisbon (Portugal) and Stykkishólmur/Reykjavík (Iceland) averaged over December–March (DJFM) for each winter.

Because cluster trajectories were smoothed to 100 normalized steps, the 22 annual DJFM NAO values (2002–2023) were linearly interpolated to 100 points using the *approx* function in *R* to enable comparison on the same relative temporal scale. For each cluster centroid, we computed Pearson's correlation coefficient (r) between the standardized cluster trajectory and the interpolated NAO series. Significance was assessed using two-tailed tests ($\alpha = 0.05$).

Overall, the correlations were mixed. For WHPT, positive associations occurred for W1 ($r = 0.34$, $p = 0.001$), W2 ($r = 0.25$, $p = 0.010$), and W4 ($r = 0.41$, $p < 0.001$), while W6 showed a negative relationship ($r = -0.25$, $p = 0.011$). For abundance and diversity both, two clusters were positively correlated with NAO (A1, A3, D1 and D3) and three negatively (A4 – A6 & D4 – D6) (**Table S6**).

### References

1. Hyndman, R.J., Athanasopoulos, G., Bergmeir, C., Caceres, G., Chhay, L., O'Hara-Wild, M., Petropoulos, F., Razbash, S. and Wang, E., 2020. Package 'forecast'.
2. Aghabozorgi, S., Shirkhorshidi, A.S. and Wah, T.Y. (2015) 'Time-series clustering – a decade review', *Information Systems*, 53, pp.16–38.
3. Pavoine, S., Love, M.S. and Bonsall, M.B., 2009. Hierarchical partitioning of evolutionary and ecological patterns in the organization of phylogenetically-structured species assemblages: application to rockfish (genus: Sebastes) in the Southern California Bight. *Ecology letters*, *12*(9), pp.898-908.
4. Aggarwal, C.C. and Reddy, C.K., 2014. Data clustering. *Algorithms and applications. Chapman&Hall/CRC Data mining and Knowledge Discovery series, Londra*.
5. Sakoe, H. and Chiba, S., 2003. Dynamic programming algorithm optimization for spoken word recognition. *IEEE transactions on acoustics, speech, and signal processing*, *26*(1), pp.43-49.
6. Rakthanmanon, T., 2012. *Efficient Algorithms for High Dimensional Data Mining*. University of California, Riverside.
7. Paparrizos, J. and Gravano, L., 2015, May. k-shape: Efficient and accurate clustering of time series. In *Proceedings of the 2015 ACM SIGMOD international conference on management of data* (pp. 1855-1870).
8. Nadaraya, E.A. (1964) 'On estimating regression', *Theory of Probability & Its Applications*, 9(1), pp.141–142.
9. Watson, G.S. (1964) 'Smooth regression analysis', *Sankhyā: The Indian Journal of Statistics, Series A*, 26(4), pp.359–372.
10. Sardá-Espinosa, A. (2017) 'Comparing time-series clustering algorithms in R using the dtwclust package', *R package vignette*, 12, pp.41.
11. Sarda-Espinosa, A., Sarda, M.A. and R, T.L.D. (2018) *dtwclust: Time series clustering along with optimisations for DTW*. R package.
12. Arbelaitz, O., Gurrutxaga, I., Muguerza, J., Pérez, J.M. and Perona, I. (2013) 'An extensive comparative study of cluster validity indices', *Pattern Recognition*, 46(1), pp.243–256.
13. Kim, M. and Ramakrishna, R.S. (2005) 'New indices for cluster validity assessment', *Pattern Recognition Letters*, 26(15), pp.2353–2363.
14. Larsen, S., Joyce, F., Vaughan, I.P., Durance, I., Walter, J.A. and Ormerod, S.J., 2024. Climatic effects on the synchrony and stability of temperate headwater invertebrates over four decades. *Global Change Biology*, *30*(1), p.e17017.
15. Moritz, S. and Bartz-Beielstein, T. (2017) 'imputeTS: Time-series missing value imputation in R', *R Journal*, 9(1), pp.207–218.

16. Bell, V.A., Kay, A.L., Rudd, A.C. and Davies, H.N., 2018. The MaRIUS-G2G datasets: Grid-to-Grid model estimates of flow and soil moisture for Great Britain using observed and climate model driving data. *Geoscience Data Journal*, *5*(2), pp.63-72.
17. Bell, V.A., Naden, P.S., Tipping, E., Davies, H.N., Carnell, E., Davies, J.A.C., Dore, A.J., Dragosits, U., Lapworth, D.J., Muhammed, S.E. and Quinton, J.N., 2021. Long term simulations of macronutrients (C, N and P) in UK freshwaters. *Science of the Total Environment*, *776*, p.145813.

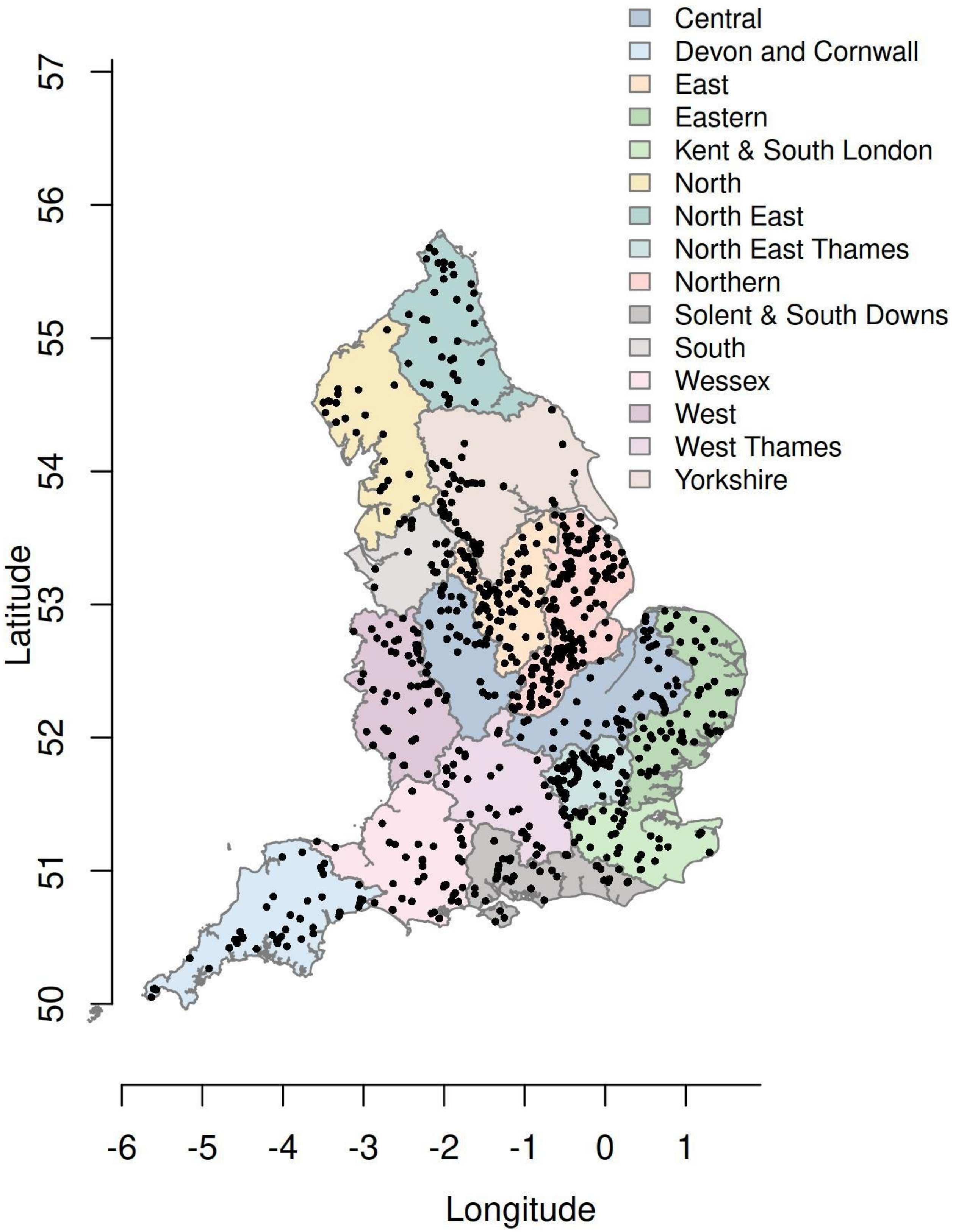

**Figure S1. Distribution of the sites from which time-series are included for unsupervised clustering.** We used a set of filtering criteria for including high resolution sites in our analysis (see main text for details). Based on these, the final dataset included 808 time-series distributed across 19 agency areas, 240 catchments, and 479 water bodies as shown above.

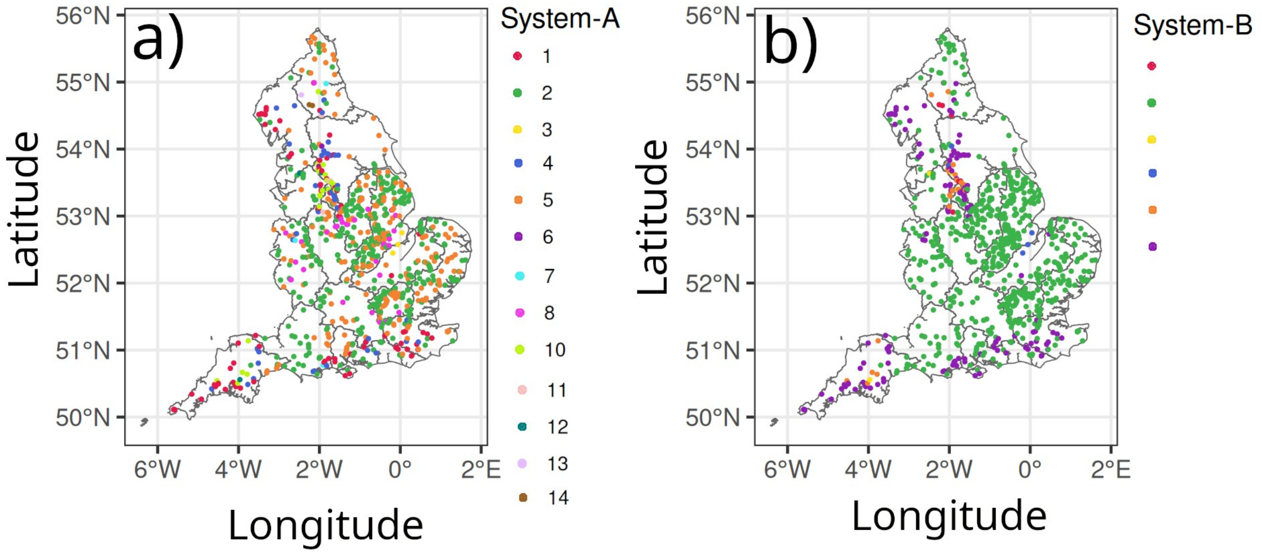

**Figure S2. Freshwater monitoring locations used in this study.** Each point represents a site, coloured according to its assigned Water Framework Directive (WFD) typology. The left panel shows typology classification following System A, whereas the right panel displays the same sites classified under System B.

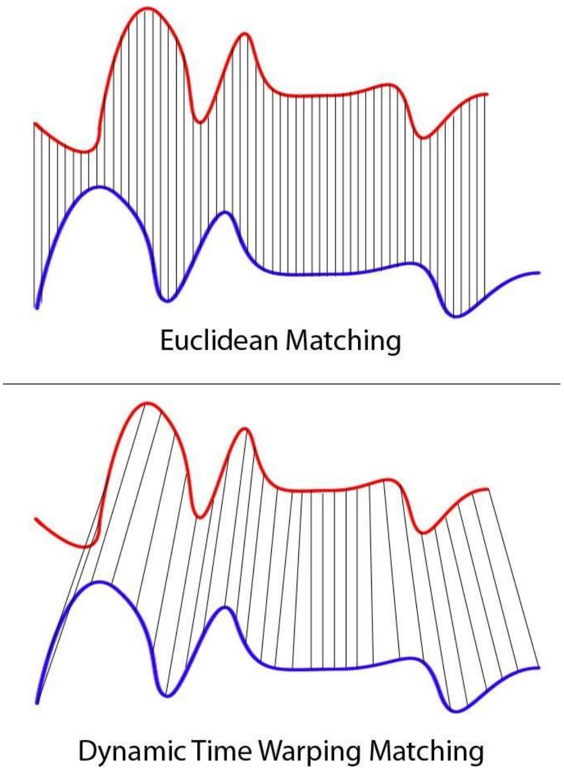

**Figure S3. Difference in matching between Euclidean and Dynamic Time Warping.** In DTW distances, two-time-series (the base time-series and new time-series) are considered similar when it is possible to map with function f(x) according to the following rules so as to match the magnitudes using an optimal (warping) path: $f(x_i)$ maps to $f(x_j)$ when $i <= j$, and $f(x_i)$ maps to $f(x_j)$ only when ($j – i$) is within a fixed pre-defined range (e.g. Sakoe-Chiba band). Image Source: Wiki Commons: File:Euclidean_vs_DTW.jpg.

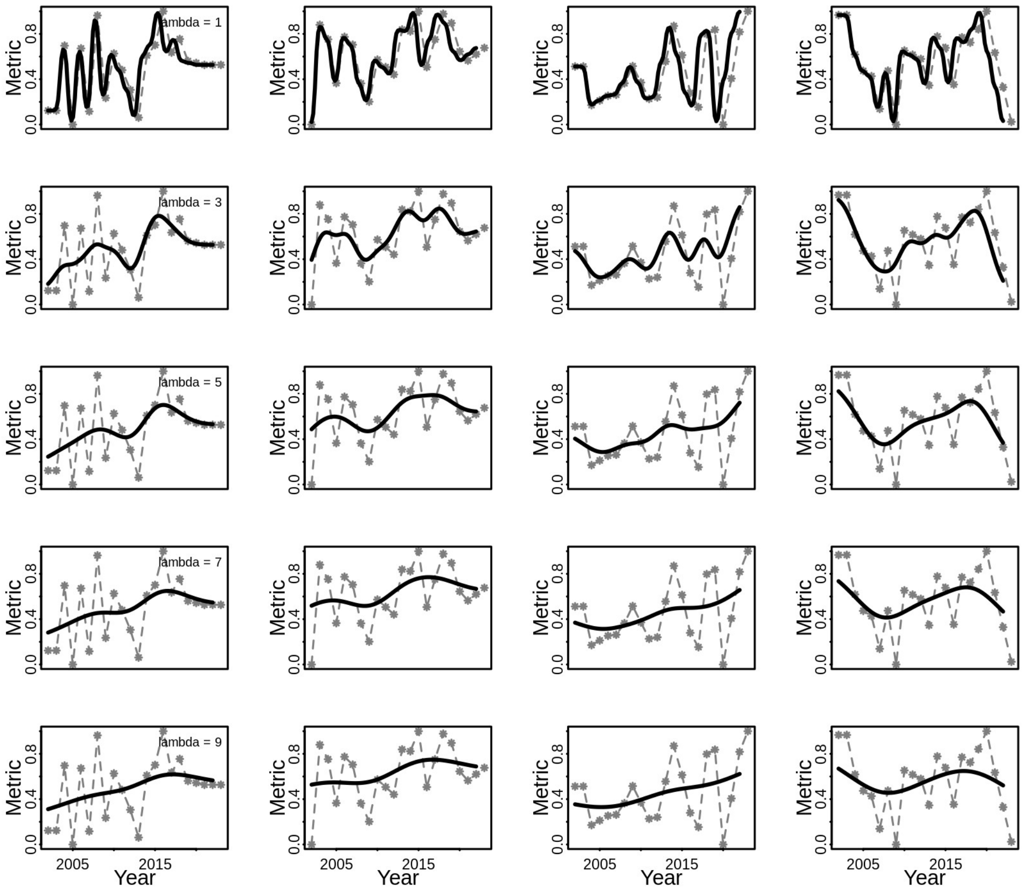

**Figure S4. Effect of the Nadaraya–Watson kernel regression smoothing parameter (λ) on raw time series.** Five different values of λ (rows) are applied to four randomly selected time series (columns) from our empirical dataset. Increasing λ results in greater smoothing. In each panel, the raw empirical time series are shown as colored lines, while the smoothed series are shown in grey.

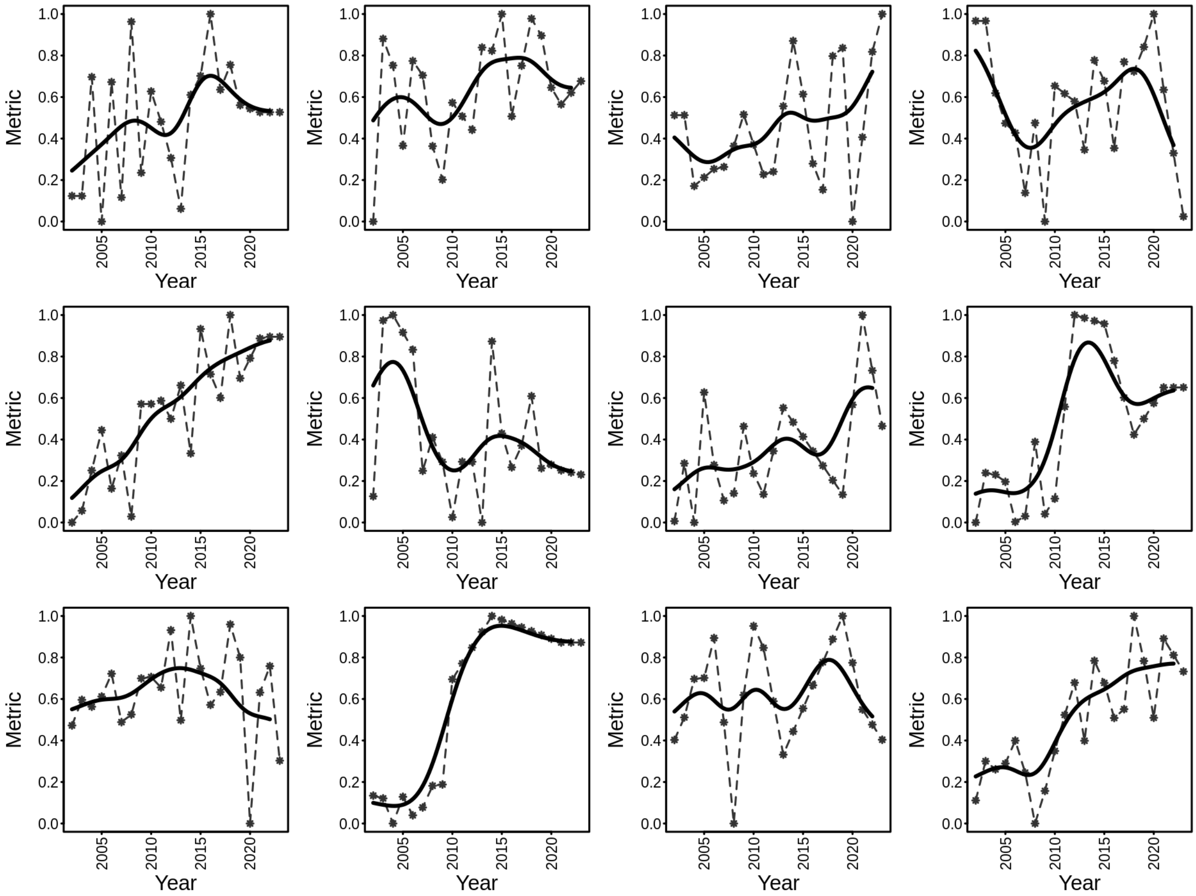

**Figure S5. Automated selection of Nadaraya–Watson kernel regression smoothing parameter lambda, based on optimization of the generalized cross validation scores.** Following a GCV optimization approach, optimal values of lambda were selected for each empirical time-series, and a random selection of 12 time-series from our empirical dataset with their smoothed counterparts are shown here. In each plot, the raw empirical time-series is shown as coloured lines and the smoothed time-series is shown in grey.

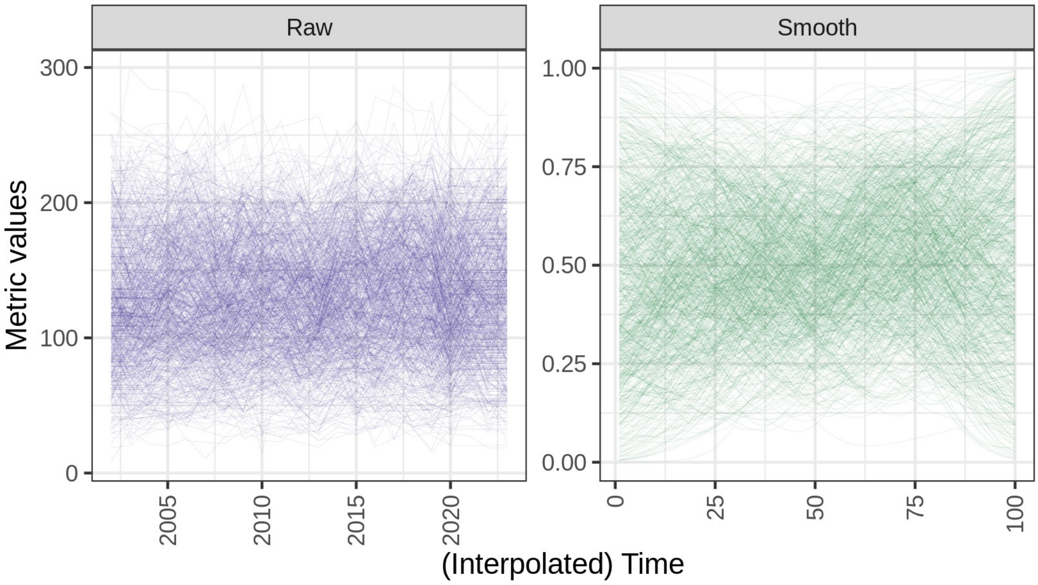

**Figure S6. Raw empirical (left panel) and smoothed (right panel) time-series for 808 WHPT metric time series.** Raw time-series are smoothed using Nadaraya–Watson kernel regression, and range normalized between 0 and 1. All clustering was performed on the smoothed, standardized dataset.

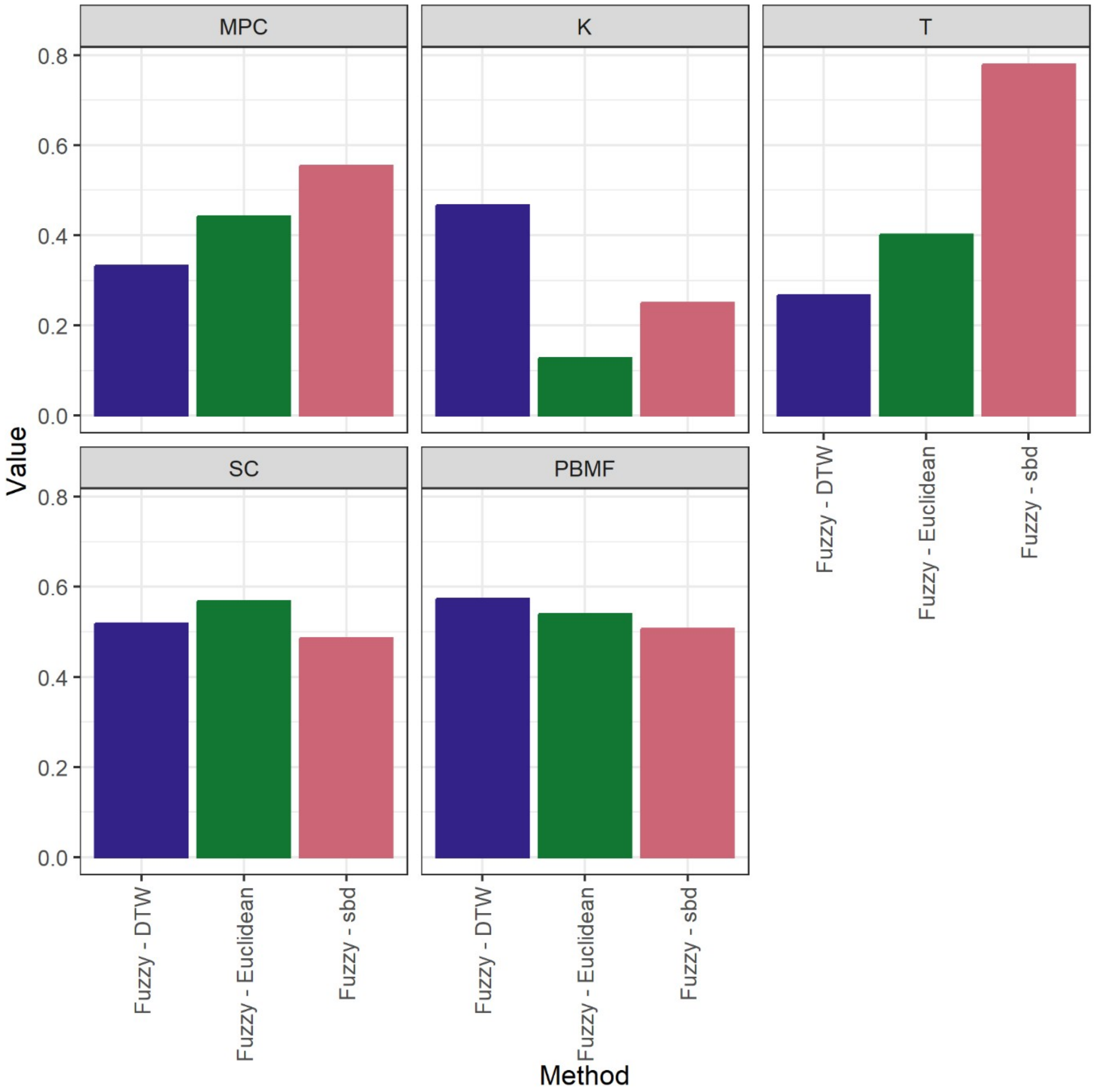

**Figure S7. Cluster Validity Index (CVI) values averaged across a range of clusters (*k* = 2 to *k* = 15) from different 'fuzzy' clustering methods.** For each *CVI*, the clustering approach with the highest average value is considered as the most efficient method. Only fuzzy indices are shown here; for crisp CVIs see Fig S7 (also see Table S3)

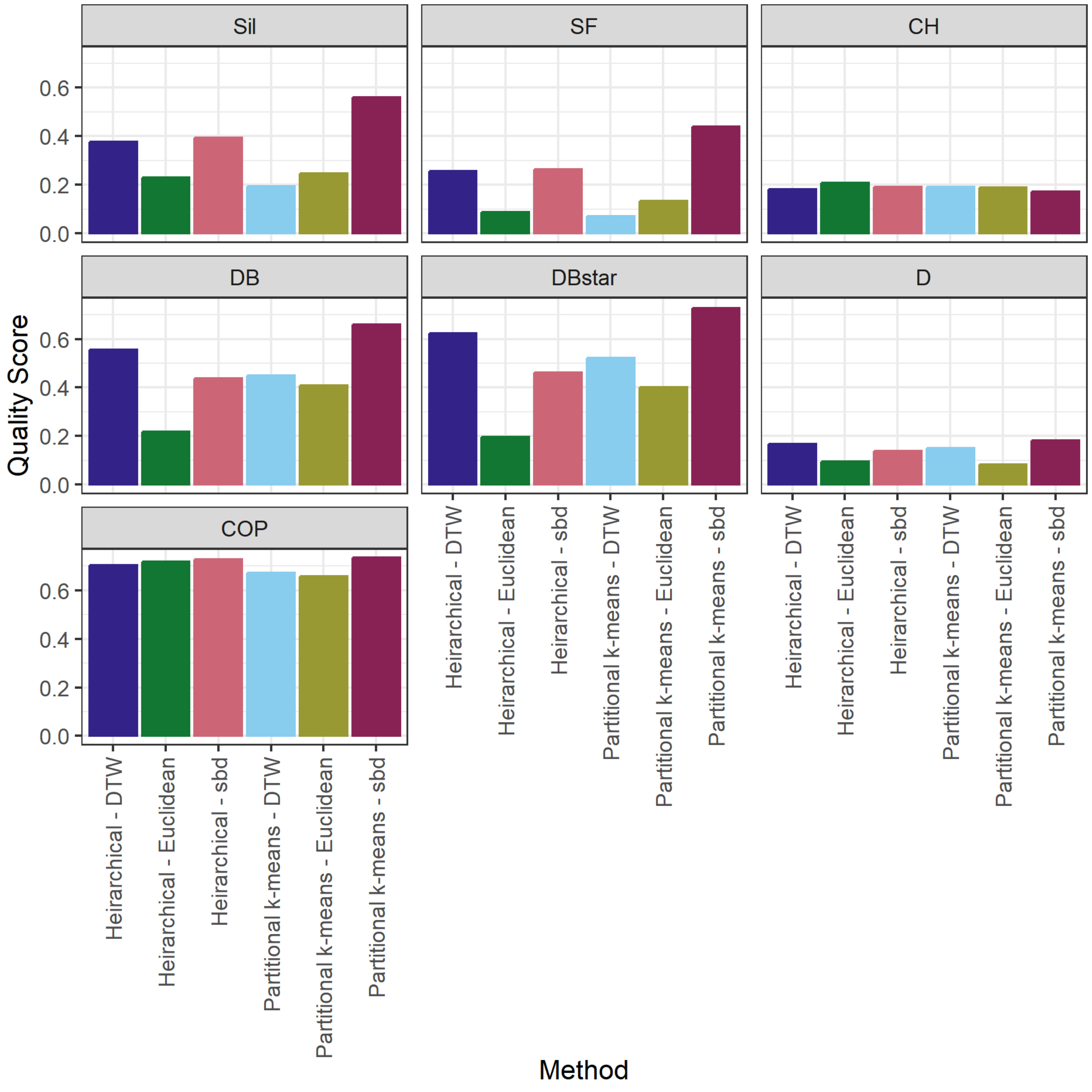

**Figure S8. Cluster Validity Index (CVI) values averaged across a range of clusters (*k* = 2 to *k* = 15) from different 'crisp' clustering methods.** For each *CVI*, the clustering approach with the highest average value is considered as the most efficient method.

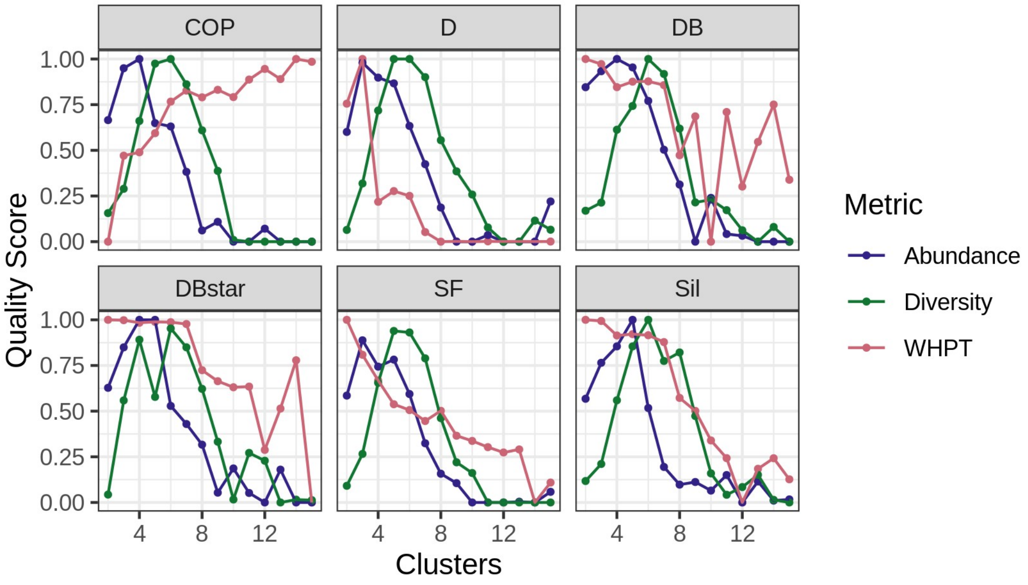

**Figure S9. Cluster Validity Index (CVI) values across a range of clusters (*k* = 2 to *k* = 15) from different clustering methods.** Each CVI is calculated iteratively for different values of *k*, and the final *k* is chosen based on (optimal) maximization of the value (see text for more details).

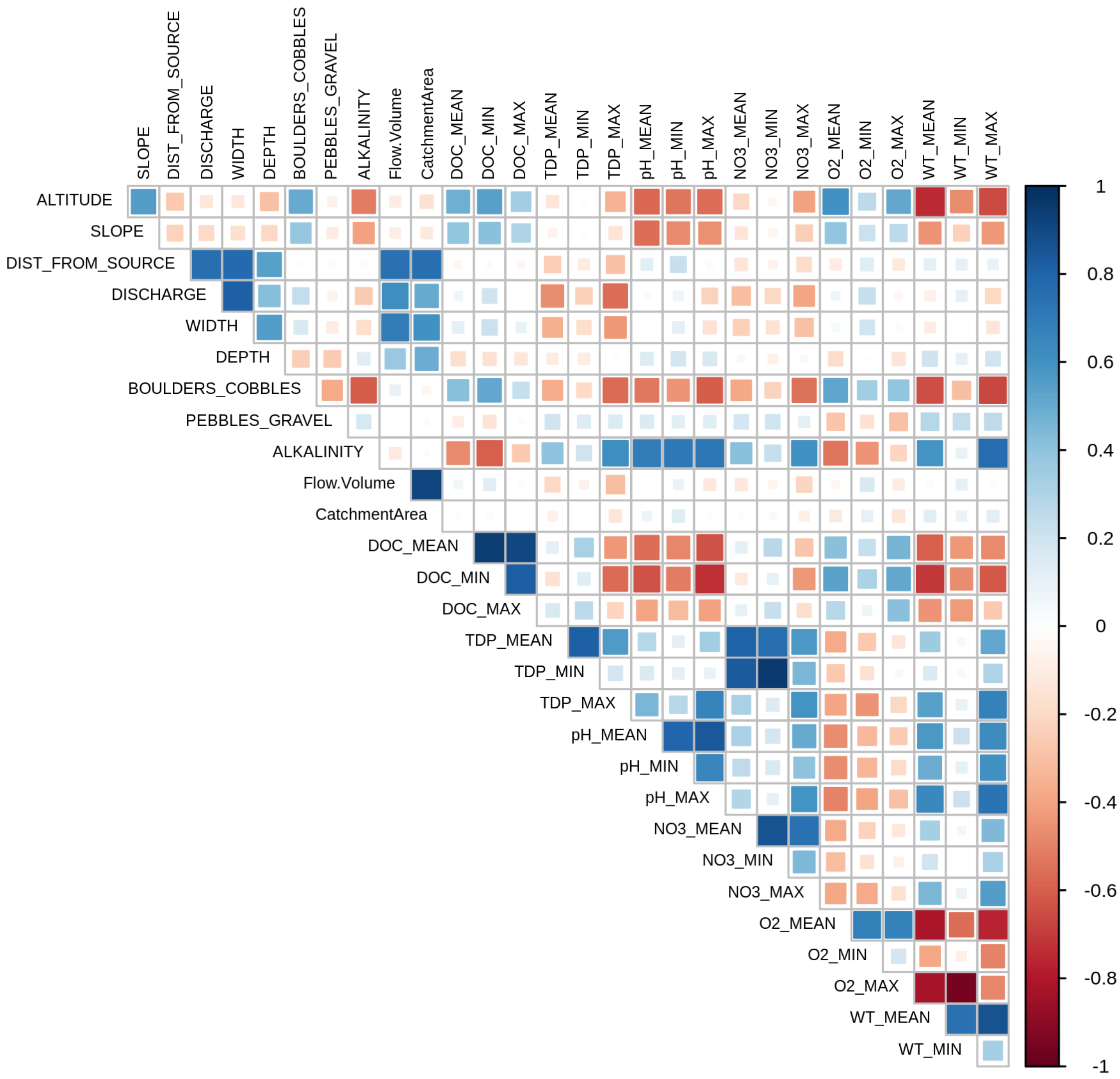

**Figure S10. Multi-collinearity between the environmental variables.** A heat map of pairwise linear regression correlation coefficients ($R^2$) among the full set of 28 environmental variables compiled for the current study. For more details, and the final subset used for all analyses, see Section S3.

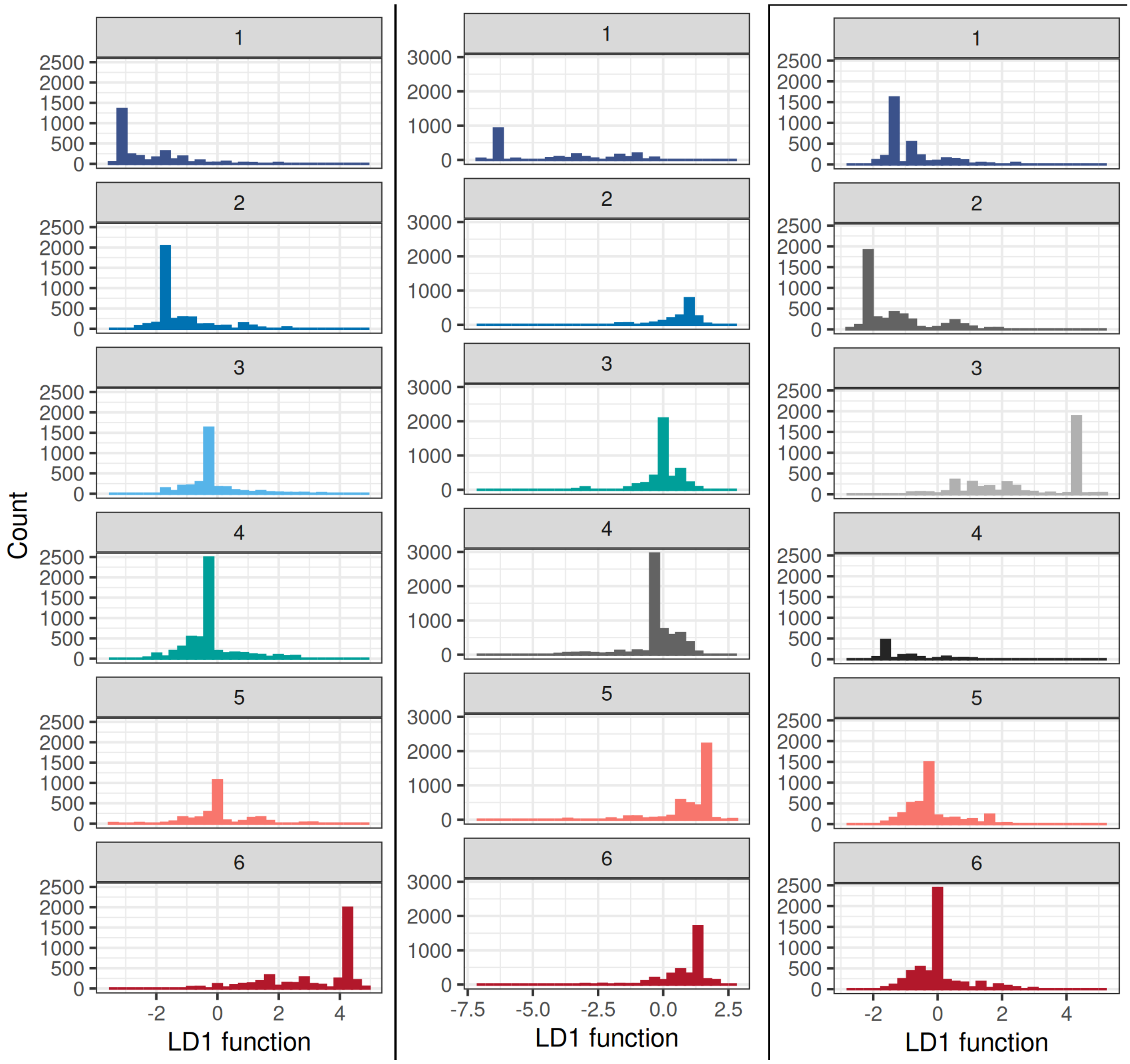

**Figure S11. Histograms of discriminant scores from Linear Discriminant Analysis (LDA), showing separation of time-series clusters for WHPT, abundance, and diversity metrics.** Each histogram displays the distribution of sites along the first linear discriminant axis, with clusters represented by distinct colours, providing a visual assessment of how well the discriminant functions derived from environmental variables distinguish among the identified time-series clusters (distribution of sites along the first two linear discriminant axis is shown in the Main Figs. 1b to 3b).

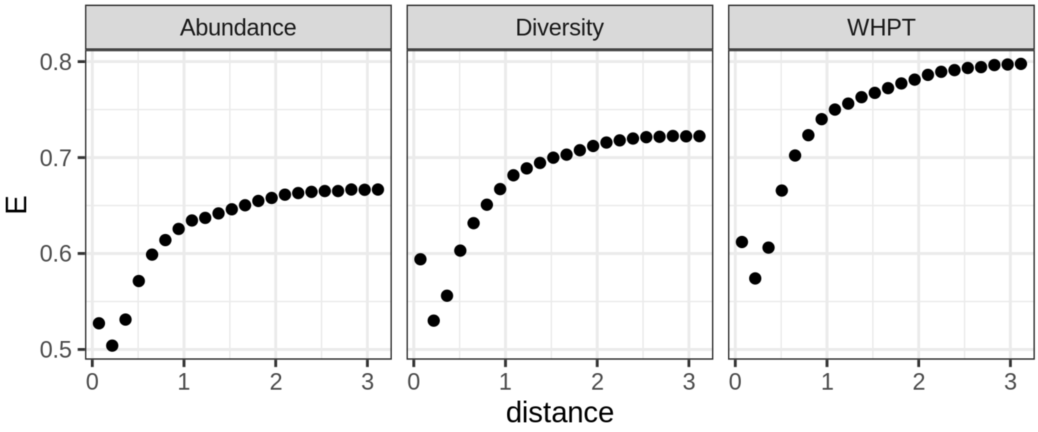

**Figure S12. Enterograms showing the global structure in spatial autocorrelation of clusters.** The enterogram E for lag distance is calculated as the mean of the ELSA statistics at different sites within the distance equal to the lag size. The visual interpretation of the graphs is that ELSA value is low within the lower distances and increases when the lag distance increases as would be expected. For the randomly distributed classes, on the other hand, this value remains at the maximum level, showing there is no spatial structure (Naimi et al. 2019). The ‘plateauing’ at around 1km suggests that maximum autocorrelation occurs at less than 1km scales in our dataset.

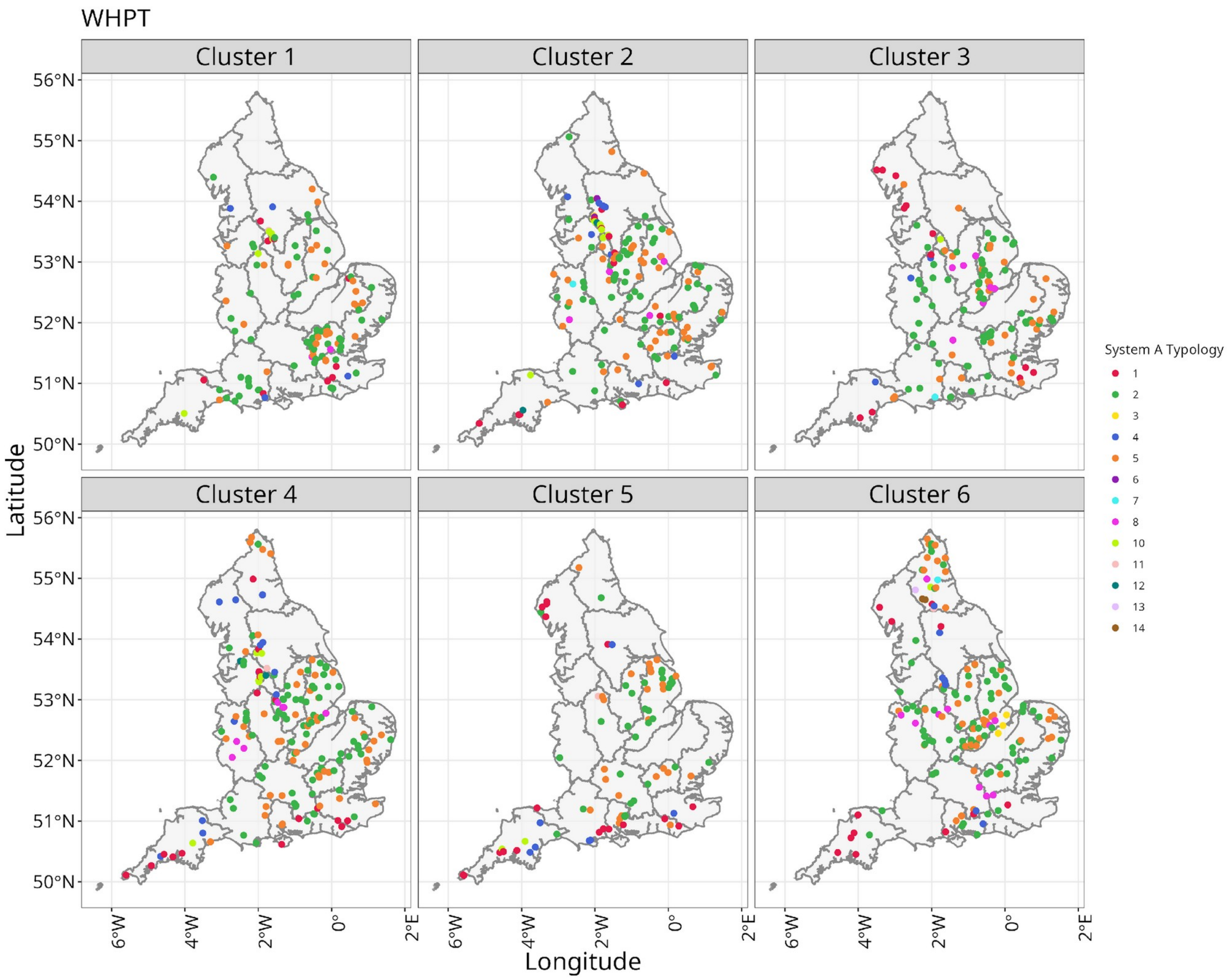

383 **Fig. S13. Spatial distribution of WHPT clusters and associated river typologies.**
384 Spatial distribution of macroinvertebrate monitoring sites assigned to the six WHPT
385 trajectory clusters. Each facet represents a distinct cluster, illustrating differences in the
386 shape of long-term pollution-sensitivity trajectories. Shaded polygons delineate
387 Environment Agency operational areas, and points indicate individual sampling sites
388 coloured by their System A typology (based on altitude, catchment size, and dominant
389 geology).

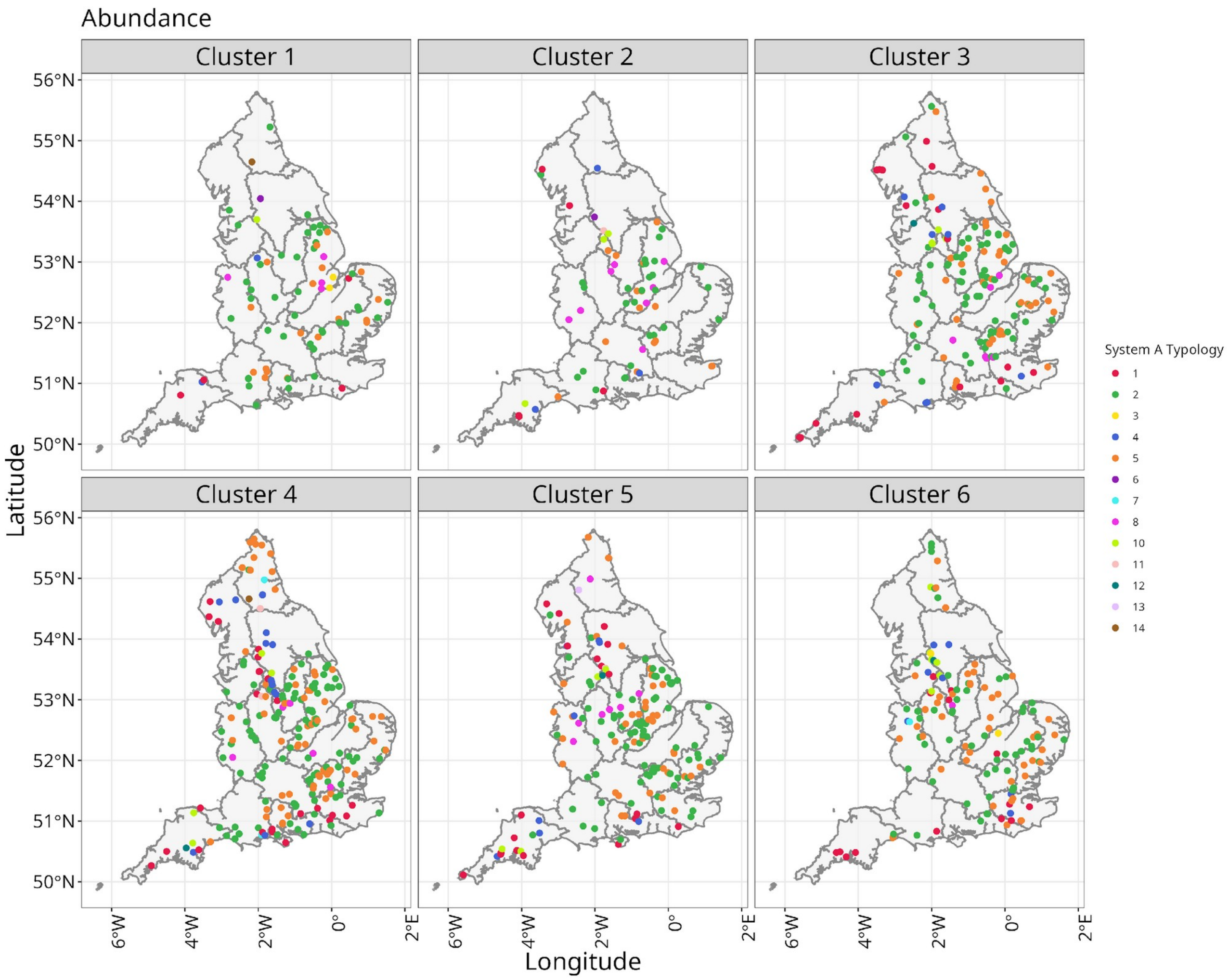

**Fig. S14. Spatial distribution of Abundance clusters and associated river typologies.** Spatial distribution of macroinvertebrate monitoring sites assigned to the six Abundance time-series clusters. Each facet represents a distinct cluster, illustrating differences in the shape of long-term macroinvertebrate temporal trends. Shaded polygons delineate Environment Agency operational areas, and points indicate individual sampling sites coloured by their System A typology (based on altitude, catchment size, and dominant geology).

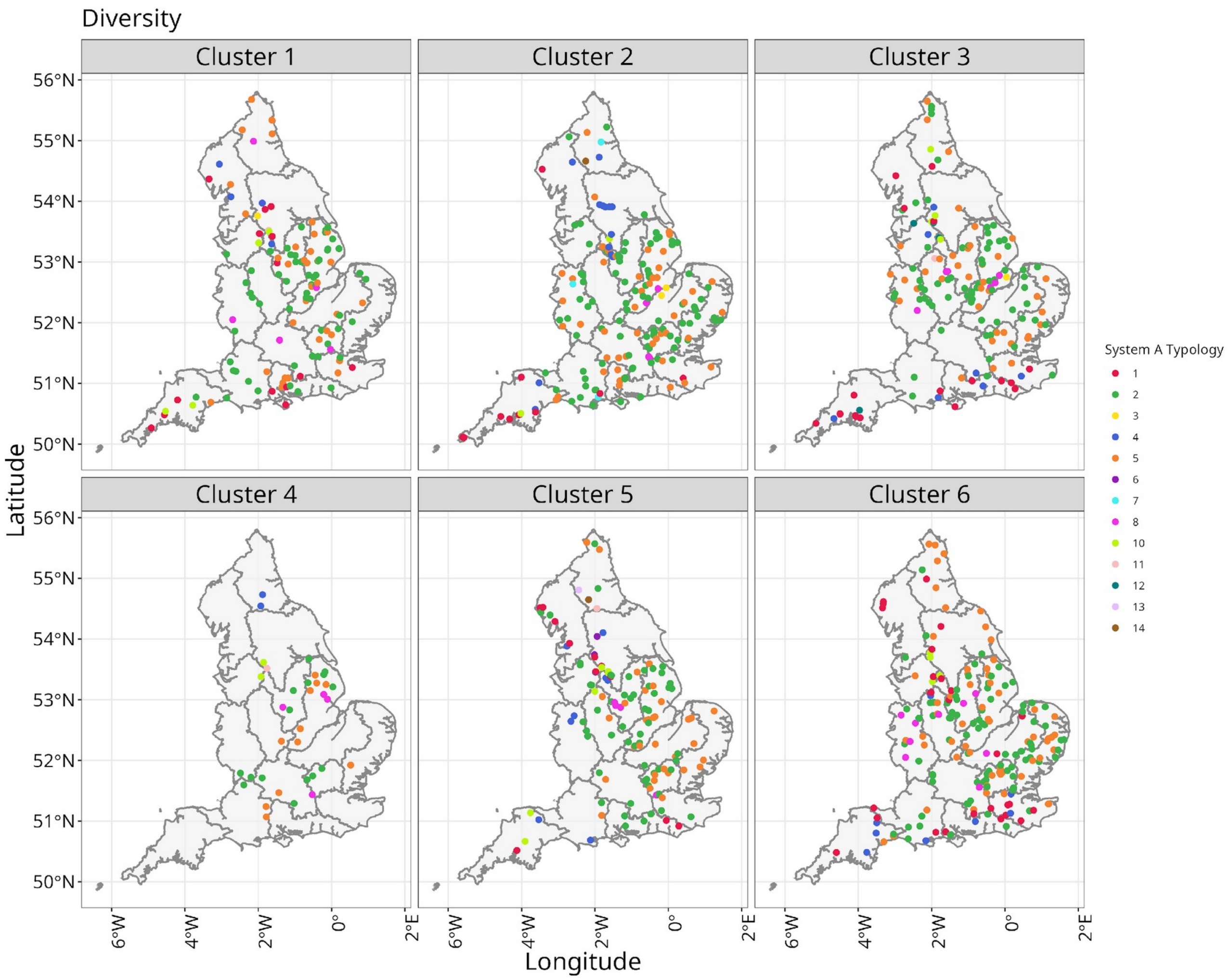

**Fig. S15. Spatial distribution of Diversity clusters and associated river typologies.** Spatial arrangement of sites classified into six clusters based on long-term diversity (Simpson's Index) trajectories. Facets correspond to the six major trajectory types, colors within each facet represent the river typology category for the different sites. Environment Agency regional boundaries are shown in grey.

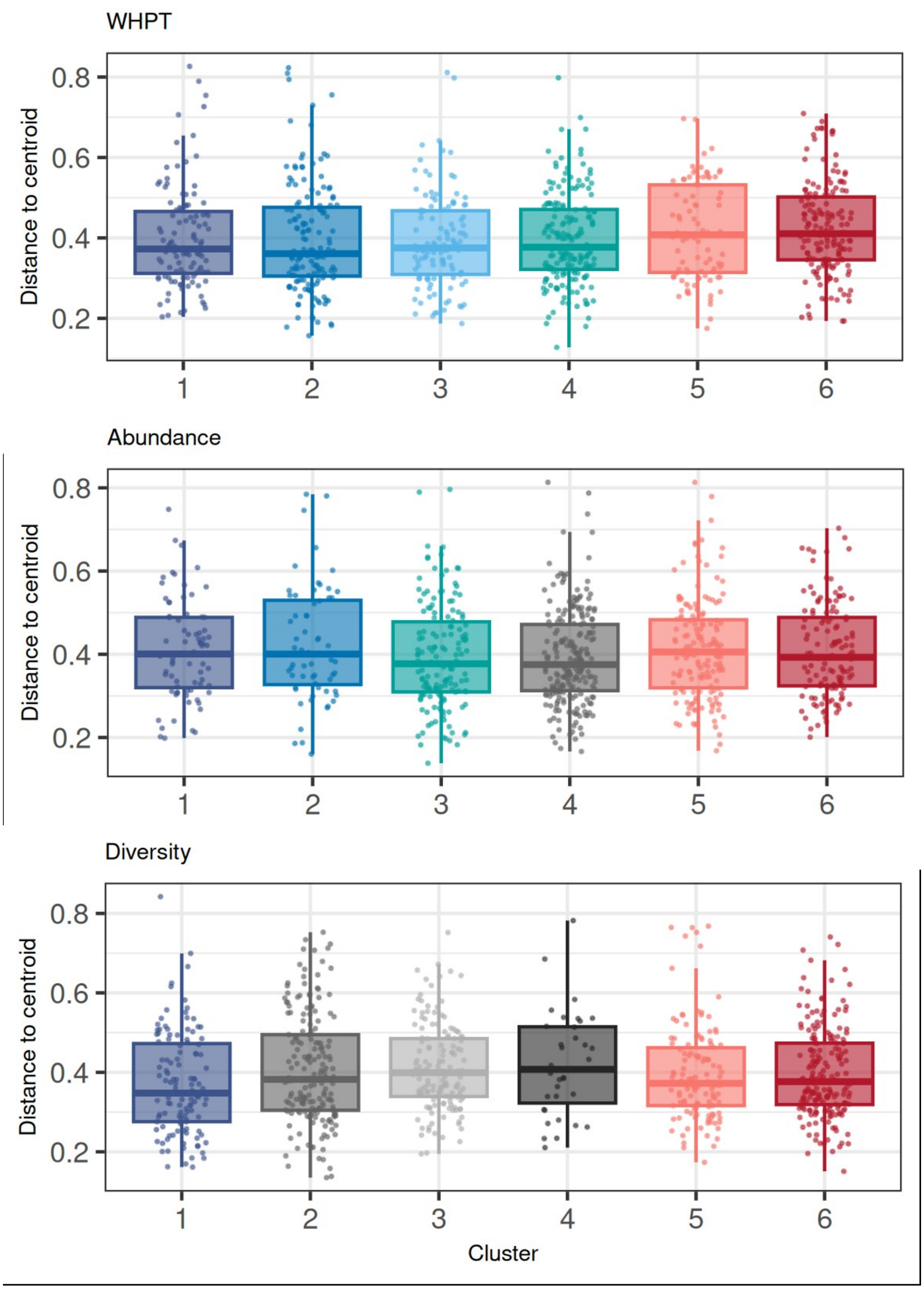

**Figure S16. Boxplots of distances to cluster centroids (multivariate dispersions) for WHPT, Abundance, and Diversity time-series clusters.** Distances represent the average Bray–Curtis dissimilarity of sites to their group centroid in ordination space, derived from the `betadisper` procedure. These plots provide a visual assessment of within-cluster variability, with wider spread or higher median distances indicating greater heterogeneity in family-level community composition. Statistical tests of homogeneity of multivariate dispersions revealed no significant differences among clusters for any of the three metrics (WHPT: $F = 1.50$, $p = 0.19$; Abundance: $F = 1.13$, $p = 0.35$; Diversity: $F = 1.91$, $p = 0.09$). Thus, the differences detected in PERMANOVA reflect true compositional shifts between clusters rather than artefacts of unequal spread.

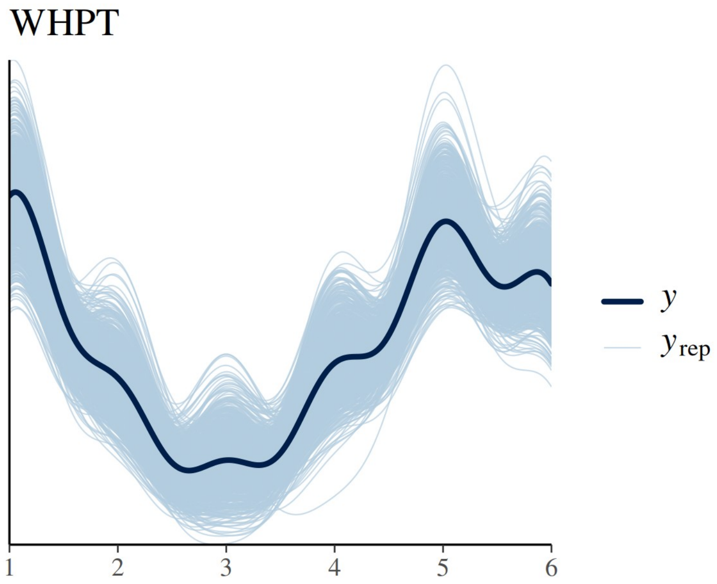

**Figure S17. Posterior predictive check for the Bayesian hierarchical model of WHPT cluster assignments.** The plot compares the observed distribution of cluster categories (dark thick blue line, *y*) with replicated data generated from the posterior predictive distribution (light blue curves, *yrep*). The x-axis represents the six categorical cluster IDs, and alignment between observed and replicated curves indicates good model fit.

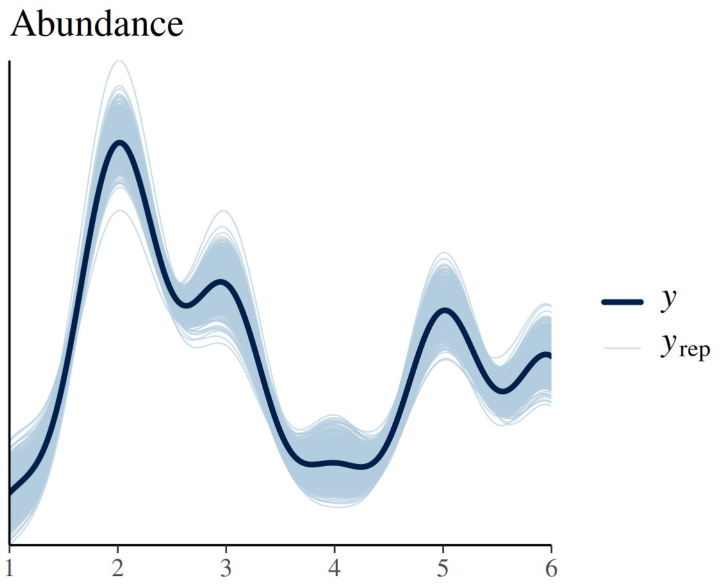

**Figure S18. Posterior predictive check for the Bayesian hierarchical model of Abundance time-series cluster assignments.**

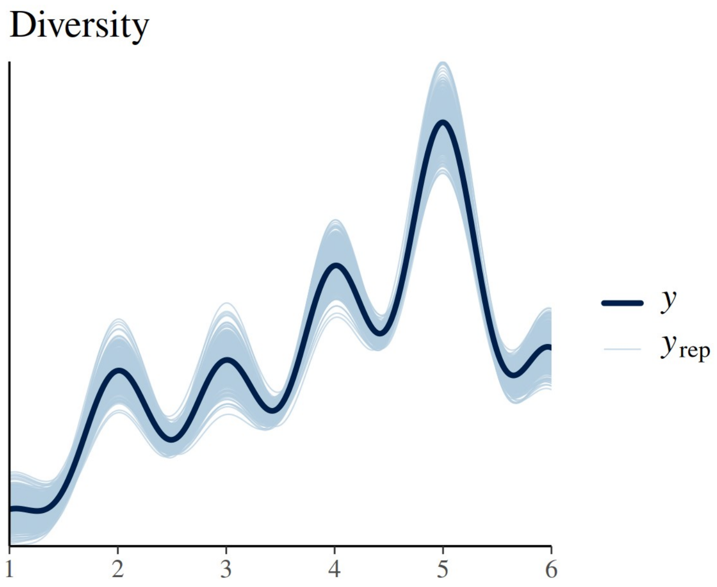

**Figure S19. Posterior predictive check for the Bayesian hierarchical model of Diversity time-series cluster assignments.**

**Table S1. Clustering methods and time-series distance measures used for unsupervised clustering of site-specific freshwater macroinvertebrate time-series based on their statistical properties.** Three different clustering methods (hierarchical, partitioning k-means, and fuzzy c-means) were combined with three distance measures: Euclidean, Dynamic Time Warping (DTW), and shape-based distance (SBD).

| Method/ Distance | Description & Equation | Strengths/Limitations in this study |
|---|---|---|
| Hierarchical (Kaufman et al. 1990) | Builds a nested hierarchy of time-series clusters by iteratively merging or splitting the clusters using a pairwise distance matrix.<br><br>$d(A,B) = \min\{d(x,y) : x \in A, y \in B\}$ | + No need to pre-define k.<br>+ Produces dendrograms useful for ecological interpretation.<br>– Sensitive to noise and outliers (issue with small datasets).<br>– Computationally expensive with many sites. |
| Partitioning k-means (MacQueen 1969; Sarda-Espinosa 2023) | Iteratively minimizes within-cluster variance.<br><br>$\text{argmin} \sum_{i=1}^{k} \sum_{x \in C_i} \|\|x - \mu_i\|\|^2$ | + Very fast and scalable (suitable for multiple sites × metrics).<br>– Requires *k* a priori.<br>– Assumes spherical clusters; may oversimplify ecological heterogeneity. |
| Fuzzy c-means (FCM; Dunn 1973; Bezdek 1981) | Assigns membership degrees $u_{ij}$ with constraint $\sum_{j=1}^{c} u_{ij}=1$.<br><br>$J_m = \sum_{i=1}^{N} \sum_{j=1}^{c} (u_{ij})^m \|\|x_i - \mu_j\|\|^2$, where m controls fuzziness. | + Captures overlapping clusters (ecologically realistic if sites share species composition patterns).<br>– Interpretation harder for large n and many clusters.<br>– Sensitive to initialization; risk of soft noise clustering. |
| Euclidean distance (Faloutsos et al. 1994) | For two series $X= (x_1,\ldots,x_t)$, $Y = (y_1,\ldots,y_t)$, $d(X,Y) = \sqrt{\sum_t (x_t - y_t)^2}$ | + Simple, fast, widely used — facilitates comparison with previous studies.<br>– Requires equal-length series; sensitive to scale and phase shifts. |
| Dynamic Time Warping (DTW; Sakoe & Chiba 1971, 1978) | Defines optimal alignment path π between X and Y:<br>$d(X,Y) = \min_\pi \sum_{(i,j) \in \pi} d(x_i, y_j)$<br>subject to monotonicity and continuity constraints. | + Allows flexible alignment, handles unequal lengths and phase shifts.<br>– Computationally intensive.<br>– Overwarping can group biologically distinct trends; Sakoe–Chiba constraint used here. |
| Shape-Based Distance (SBD; Paparrizos & Gravano 2015) | Based on normalized cross-correlation:<br>$d(X,Y) = 1 - \max_w NCC_w(X,Y)$,<br>where NCC is coefficient-normalized correlation at lag w. | + Captures shape similarity regardless of amplitude; suitable with site-specific scaling differences.<br>– Requires normalized inputs; sensitive to strong outliers.<br>– Less interpretable biologically than DTW. |

*d(A,B): distance between clusters A and B;*
*d(x,y)distance between individual time-series elements x and y;*
*k: number of clusters;*
*$\mu_i$: centroid (prototype) of cluster i;*
*$C_i$: set of time series assigned to cluster I;*
*$u_{ij}$: degree of membership of time series i in cluster j;*
*m: fuzziness parameter in fuzzy c-means (m > 1, typically m=2);*
*$J_m$: objective function minimized in fuzzy c-means clustering.*
*X, Y: two time series ($X=(x_1,\ldots,x_t)$, $Y=(y_1,\ldots,y_t)$).*

**Table S2. Cluster Validity Indices (CVIs) used to compare clustering approaches and select the optimum number of clusters.** To evaluate clustering efficiency across various time-series configurations, we employed a combination of seven crisp and five fuzzy validity indices. These indices typically measure both cluster cohesion (internal similarity) and separation (between-cluster difference), combining them into a single quality score. A brief description, along with the original reference for each CVI is listed below.

| CVI | Description |
|---|---|
| **For "Crisp" clusters (Partitional and Hierarchical)** | |
| Silhouette (Sil); higher the better; Rousseeuw, 1987 | The cohesion is measured based on the distance between all the points in the same cluster and the separation is based on the nearest neighbour distance. |
| Score Function (SF); higher the better; Saitta et al. (2007) | The separation is measured based on the distance from the cluster centroids to the global centroid and the cohesion is based on the distance from the points in a cluster to its centroid. |
| Calinski–Harabasz (CH); higher the better; Milligan & Cooper, 1985; Arbelaitz et al. (2013) | The cohesion is estimated based on the distances from the points in a cluster to its centroid. The separation is based on the distance from the centroids to the global centroid. |
| Davies–Bouldin (DB); lower the better; Davies & Bouldin, 1975; Arbelaitz et al. (2013) | This is probably one of the most used indices in CVI comparison studies. It estimates the cohesion based on the distance from the points in a cluster to its centroid and the separation based on the distance between centroids. |
| Modified Davies-Bouldin index (DBstar); lower the better; Kim and Ramakrishna (2005 | A variation of the Davies–Bouldin index |
| Dunn index (D); higher the better; Arbelaitz et al. (2013) | The cohesion is estimated by the nearest neighbour distance and the separation by the maximum cluster diameter. |
| COP; Lower the better; Gurrutxaga et al. 2010 | The cohesion is estimated by the distance from the points in a cluster to its centroid and the separation is based on the furthest neighbour distance. Called COP because it satisfies the Context-independent Optimality and Partiality properties |
| **For "overlapping" clusters (Fuzzy)** | |
| Modified Partition Coefficient (MPC); higher the better; Dave, 1996 | Based on minimizing the overall content of pairwise fuzzy intersection. It indicates the average relative amount of membership sharing done between pairs of fuzzy subsets |
| (K); higher the better; Xie and Beni (1991); Kwon 1998 | Focused on two properties: compactness and separation, and further added a punishment function to eliminate its tendency to monotonically decrease when the number of clusters approaches to the number of data points |
| (T); lower the better; Yang et al. (2005) | *ad hoc* punishing function (average distance between cluster centroids) applied to eliminate the decreasing tendency as $c \rightarrow n$, moreover, the second term in denominator is also a punishing function used to strengthen the numerical stability as $m \rightarrow \infty$. |
| (SC); higher the better; Zahid et al. 1999 | Considers the compactness and separation by analysing the geometrical properties of the data structure and membership functions, and then uses a fuzzy union and a fuzzy intersection to obtain the fuzzy compactness/fuzzy separation degree |
| (PBMF); higher the better; Pakhira et al. (2004) | Composed of three factors: divisibility of a *c* cluster system; the sum of weighted intra-cluster distances for the complete data set; and the maximum inter-cluster separation. |

**Table S3. Cluster Validity Index (CVI) values averaged across a range of clusters (k = 2 to k = 15) from different clustering methods.** For each CVI, the highest value is highlighted in bold. One index (CH) was found to be highly similar across methods and is thus considered uninformative here (for full names and further details on the indices, see **Table S2;** also see **Figs. S6-S8**).

| **Method - Crisp** | **Sil** | **SF** | **CH** | **DB** | **DBstar** | **D** | **COP** |
|---|---|---|---|---|---|---|---|
| Hierarchical - DTW | 0.377 | 0.255 | 0.183 | 0.555 | 0.624 | 0.167 | 0.703 |
| Hierarchical - Euclidean | 0.230 | 0.088 | 0.208 | 0.217 | 0.197 | 0.095 | 0.718 |
| Hierarchical - sbd | 0.395 | 0.263 | 0.190 | 0.438 | 0.462 | 0.139 | 0.726 |
| Partitional k-means - DTW | 0.194 | 0.072 | 0.192 | 0.450 | 0.521 | 0.152 | 0.672 |
| Partitional k-means - Euclidean | 0.246 | 0.134 | 0.188 | 0.408 | 0.401 | 0.084 | 0.657 |
| Partitional k-means - sbd | **0.560** | **0.439** | 0.173 | **0.660** | **0.726** | **0.183** | **0.733** |
| **Method - Fuzzy** | **MPC** | **K** | **T** | **SC** | **PBMF** | | |
| Fuzzy - DTW | 0.33 | 0.47 | 0.27 | 0.52 | 0.53 | | |
| Fuzzy - Euclidean | 0.44 | 0.13 | 0.40 | 0.57 | 0.54 | | |
| Fuzzy - sbd | 0.56 | 0.25 | 0.78 | 0.49 | 0.51 | | |

**Table S4. Proportion of trace explained by each linear discriminant (LD) axis for WHPT, abundance, and diversity models.** In linear discriminant analysis (LDA), the *proportion of trace* represents the fraction of between-group variance explained by each discriminant function. Each LD axis is an orthogonal linear combination of predictor variables that maximizes group separation, with LD1 explaining the greatest share of discrimination, followed by subsequent axes. The values are computed as the squared singular values for each LD axis divided by the sum across all axes, such that the proportions sum to 1. Higher values indicate that the corresponding axis captures more of the discriminatory signal among clusters. For example, LD1 explains ~29% of the variance in WHPT, and ~25% in abundance and diversity, with the remaining variance distributed across the other discriminant axes.

| | **Proportion of Trace** | | | | |
|---|---|---|---|---|---|
| | **LD1** | **LD2** | **LD3** | **LD4** | **LD5** |
| **WHPT** | 0.29 | 0.22 | 0.19 | 0.17 | 0.13 |
| **Abundance** | 0.25 | 0.22 | 0.20 | 0.17 | 0.16 |
| **Diversity** | 0.25 | 0.21 | 0.20 | 0.18 | 0.16 |

**Table S5. Variable importance for WHPT, abundance, and diversity linear discriminant models.** For each model (WHPT, abundance, diversity), the standardized coefficients for each predictor variable were extracted from the corresponding LDA model. The contribution of each predictor to discrimination among clusters was quantified as the weighted sum of its absolute coefficients across all discriminant functions (LD1 to LD5), where the weights were the proportion of variance explained by each linear discriminant axis (i.e., squared singular values divided by the total, following Wilks 1932; Fisher 1936; see also Venables & Ripley 2002). The resulting variable scores were normalized to percentages by dividing each variable's contribution by the total summed contribution across all predictors, so that the final values reflect the relative importance (%) of each variable in separating clusters. The top three most contributing variables foreach metric are highlighted in shades of grey.

| | **Importance** | | |
|---|---|---|---|
| **Variable** | **WHPT** | **Abundance** | **Diversity** |
| pH_MAX | 18.00 | 16.96 | 12.83 |
| TDP_MAX | 13.33 | 15.42 | 16.67 |
| DIST_FROM_SOURCE | 11.20 | 9.18 | 15.21 |
| pH_MIN | 10.13 | 18.26 | 8.92 |
| ALTITUDE | 9.98 | 4.00 | 8.89 |
| SLOPE | 9.50 | 6.78 | 8.30 |
| NO3_MAX | 7.35 | 9.07 | 6.48 |
| BOULDERS_COBBLES | 5.59 | 6.94 | 4.97 |
| O2_MIN | 5.44 | 6.37 | 4.94 |
| PEBBLES_GRAVEL | 5.24 | 2.77 | 4.20 |
| DEPTH | 4.23 | 4.26 | 8.58 |

**Table S6.** Pearson correlation between smoothed cluster centroids and the winter North Atlantic Oscillation (NAO; DJFM) index (2002–2023). NAO values were linearly interpolated to a 100-point normalized time axis to match cluster trajectories.

| Metric | Cluster | r | p.value |
|---|---|---|---|
| Abundance | Cluster_1 | 0.24 | 0.016 |
| | Cluster_2 | 0.09 | 0.359 |
| | Cluster_3 | 0.43 | 0.000 |
| | Cluster_4 | -0.25 | 0.013 |
| | Cluster_5 | -0.20 | 0.044 |
| | Cluster_6 | -0.26 | 0.010 |
| Diversity | Cluster_1 | 0.27 | 0.007 |
| | Cluster_2 | -0.19 | 0.058 |
| | Cluster_3 | 0.29 | 0.003 |
| | Cluster_4 | -0.41 | 0.000 |
| | Cluster_5 | -0.20 | 0.046 |
| | Cluster_6 | -0.27 | 0.007 |
| WHPT | Cluster_1 | 0.34 | 0.001 |
| | Cluster_2 | 0.25 | 0.010 |
| | Cluster_3 | 0.08 | 0.410 |
| | Cluster_4 | 0.41 | 0.000 |
| | Cluster_5 | -0.13 | 0.203 |
| | Cluster_6 | -0.25 | 0.011 |
